# Single-cell lung eQTL dataset of Asian never-smokers highlights the roles of alveolar cells in lung cancer etiology

**DOI:** 10.64898/2026.03.26.714500

**Authors:** Thong Luong, Jinhu Yin, Bolun Li, Ju Hye Shin, Elelta Sisay, Sama Mikhail, Fei Qin, Samuel Anyaso-Samuel, Alexander Kane, Alyxandra Golden, Jia Liu, Chia Han Lee, Zixuan Eleanor Zhang, Yoon Soo Chang, Jinyoung Byun, Younghun Han, Maria Teresa Landi, Nicholas Mancuso, Nicholas E. Banovich, Nathaniel Rothman, Christopher Amos, Qing Lan, Kai Yu, Tongwu Zhang, Erping Long, Jianxin Shi, Jin Gu Lee, Eun Young Kim, Jiyeon Choi

## Abstract

Single-cell expression quantitative trait loci (sc-eQTL) analyses are powerful in identifying context-specific susceptibility genes from genome-wide association studies (GWAS) loci. However, few studies have comprehensively investigated cells of lung cancer origin in non-European populations. Here, we built a lung sc-eQTL dataset from 129 Korean women never-smokers with epithelial cell enrichment. eQTL mapping identified 2,229 genes with an eQTL in 33 cell types, including East Asian-specific findings when compared to predominantly European datasets. Integration with single-cell chromatin accessibility data demonstrated an enrichment of cell-type specific eQTLs in cell-type matched candidate enhancers, while shared eQTLs were more frequently found near promoters. Colocalization and transcriptome-wide association study unveiled 36 susceptibility genes from 22 cell types in 22 lung cancer loci, including 10 loci not achieving genome-wide significance in prior GWAS. Around 47% of these genes were from cells of the alveoli, underscoring their importance, especially in lung adenocarcinoma (LUAD) susceptibility. Focusing on the trajectory of alveolar epithelial cell regeneration, we detected 785 cell-state-interacting QTLs, which overlapped with 28% (10) of the identified susceptibility genes. Finally, we experimentally validated East Asian-and alveolar type 2 cell-specific eQTL of *TCF7L2* underlying East Asian LUAD locus, 10q25.2. Consistent with its role as a Wnt/β-catenin effector, TCF7L2 displayed significant effect on lung adenocarcinoma cell growth. Our data highlighted context-specific susceptibility genes, especially from alveolar cells of lung, contributing to lung cancer etiology.

## Introduction

World-wide, lung cancer is one of the most common and deadly forms of cancer, and the prevalence can differ among populations by histological subtypes, smoking status, geographical location or ancestry, and sex^1^. For example, lung adenocarcinoma (LUAD) accounts for >40% of all lung cancer cases, wherein incidence rates among never-smokers are higher in women, especially those of Asian descent^2–4^. Although smoking is the main risk factor, up to 25% of lung cancer cases arise in never-smokers^5^. Genetic factors also contribute to lung cancer risk in both smokers and never-smokers, and heritability is estimated to be around 15-18%^6–10^.

Consistently, genome wide association studies (GWAS) have identified multiple genomic loci associated with lung cancer risk, including those specific to smoking status, histological subtypes, and population^11–14^. However, how these variants contribute to lung cancer susceptibility in different contexts is still not well understood, as they predominantly reside in non-protein-coding regions and are hypothesized to contribute to risk mainly by altering regulatory elements and subsequent gene expression. Expression quantitative trait loci (eQTL) analysis links variants to gene expression changes, enabling nomination of lung cancer susceptibility genes^15^. Although powerful, these eQTL studies were traditionally based on bulk tissues using average gene expression across different cell types and do not fully explain the heritability underlying trait-associated loci^16^. For example, a recent transcriptome-wide association study (TWAS) using bulk lung tissue eQTL (n=1,038 individuals) only identified genes for 17 out of 45 lung cancer risk loci^17^. Apart from non-transcriptional mechanisms not explained by eQTL, a potential reason for a low discovery rate is that candidate causal variants (CCVs) and susceptibility genes function in a cell type and context-dependent manner, necessitating more context-specific datasets to delineate their roles in disease etiology^18^.

To this end, single-cell RNA sequencing (scRNA-seq) has revealed an astonishing complexity of lung cell types, with more than 60 reported cell types, including heterogeneous cell states, such as progenitor and transitional cell types (e.g., “AT0” and “RAS”)^19–22^. These progenitor or transitional cells alongside well-known facultative progenitor, alveolar type 2, or AT2, are suggested to be involved in maintaining the lung alveoli and regenerating it following injury, thereby contributing to chronic lung diseases such as chronic obstructive pulmonary disease (COPD) and lung adenocarcinoma^20,21,23^. scRNA-seq has also facilitated single-cell eQTL (sc-eQTL) mapping, which have been instrumental in detecting context-specific genetic regulation underlying GWAS loci for common diseases^24–27^. However, tissue based sc-eQTL datasets are still relatively scarce, due to the logistical and financial constraints of building a large scale scRNA-seq datasets. Moreover, in the context of solid cancer, epithelial cells, where most cancers originate, are often depleted during typical scRNA-seq sample processing (i.e., tissue dissociation and freeze-thaw process), whereas heartier immune cells persist. Despite these constraints, sc-eQTL datasets from lung tissues have emerged, and were able to unveil contexts underlying genetic susceptibility to chronic lung diseases or lung cancer even at a smaller sample size than previous bulk eQTL studies^26,28^. However, datasets from non-European populations remain scarce. For example, a recent study from Natri et al (n=113 individuals) is mainly of European population (66.7% of the dataset) and highlighted disease-normal comparisons (57.9% of participants with Interstitial Lung Disease or ILD)^26^. A recent Chinese dataset from Fu et al (n=222 individuals) identified promising candidate susceptibility genes for Chinese non-small cell lung cancer GWAS^28^. However, susceptibility genes for many loci are still not identified, and assessments of East Asian ancestry-specific sc-eQTL findings are largely lacking.

To enable characterization of lung cancer GWAS in a diverse population while minimizing confounding effects of smoking and sex in a modest sample size, we built a sc-eQTL dataset of tumor-distant normal lung tissues from 129 Korean women who were never-smokers. To enrich and preserve epithelial cells, we utilized fluorescence activated cell sorting (FACS) and cell type balancing. Our eQTL mapping in 33 cell types revealed that shared or cell type and lineage specific eQTLs aligned with relevant functional elements. Integration of lung cancer GWAS summary statistics with sc-eQTL provided context to known susceptibility genes, while nominating novel genes and loci. Lastly, experimental validation of unique susceptibility genes from Asian never-smoker GWAS loci provided insights into lung adenocarcinoma etiology.

Overall, our sc-eQTL dataset increases the diversity in current genomic resources and enables us to uncover context-specific genetic regulation and lung cancer susceptibility genes.

## Results

### Epithelial cell-enriched single-cell dataset represents cells of lung cancer origin

To build a lung sc-eQTL dataset from 129 Korean women never-smokers, we utilized a multiplexing and cell type balancing strategy (Fig. 1). Our design was focused on minimizing sample heterogeneity by using tumor-distant normal lung tissues of all female never-smokers from a single center and standardizing the process. All patients were treatment-naïve at the time of surgery, mainly for early stage LUAD (∼79% stage I) (Baseline characteristics in Supplementary Table 1). Although lung tumors predominantly arise from an epithelial cell type, epithelial cells are often underrepresented in single cell capture while immune cells are preferentially retained^19,29^. To alleviate this issue, we utilized FACS and cell type balancing as previously described^29^ (Fig. 1a, Methods). Balanced cells were multiplexed with ∼6 samples per batch for single cell capture and sequencing. The number of individuals per batch was optimized to reduce doublets while maximizing the number of individuals with sufficient cells for eQTL detection power^30^ (Supplementary Fig. 1). Balancing of cell types and multiplexing further allowed uniform processing of samples while addressing sampling bias of cell type proportions. After de-multiplexing and quality control, we detected 360,127 lung cells from 129 individuals assayed in 23 batches (Supplementary Table 2-3). Consistent with the initial power calculation, a median of 2,770 cells/individuals (range = 340 - 8,036) were retained^30^ (Supplementary Tables 3-5). This enabled us to perform sc-eQTL mapping and context-specific lung cancer susceptibility genes nomination, followed by functional characterization (Figs. 1b-d).

**Figure 1.**
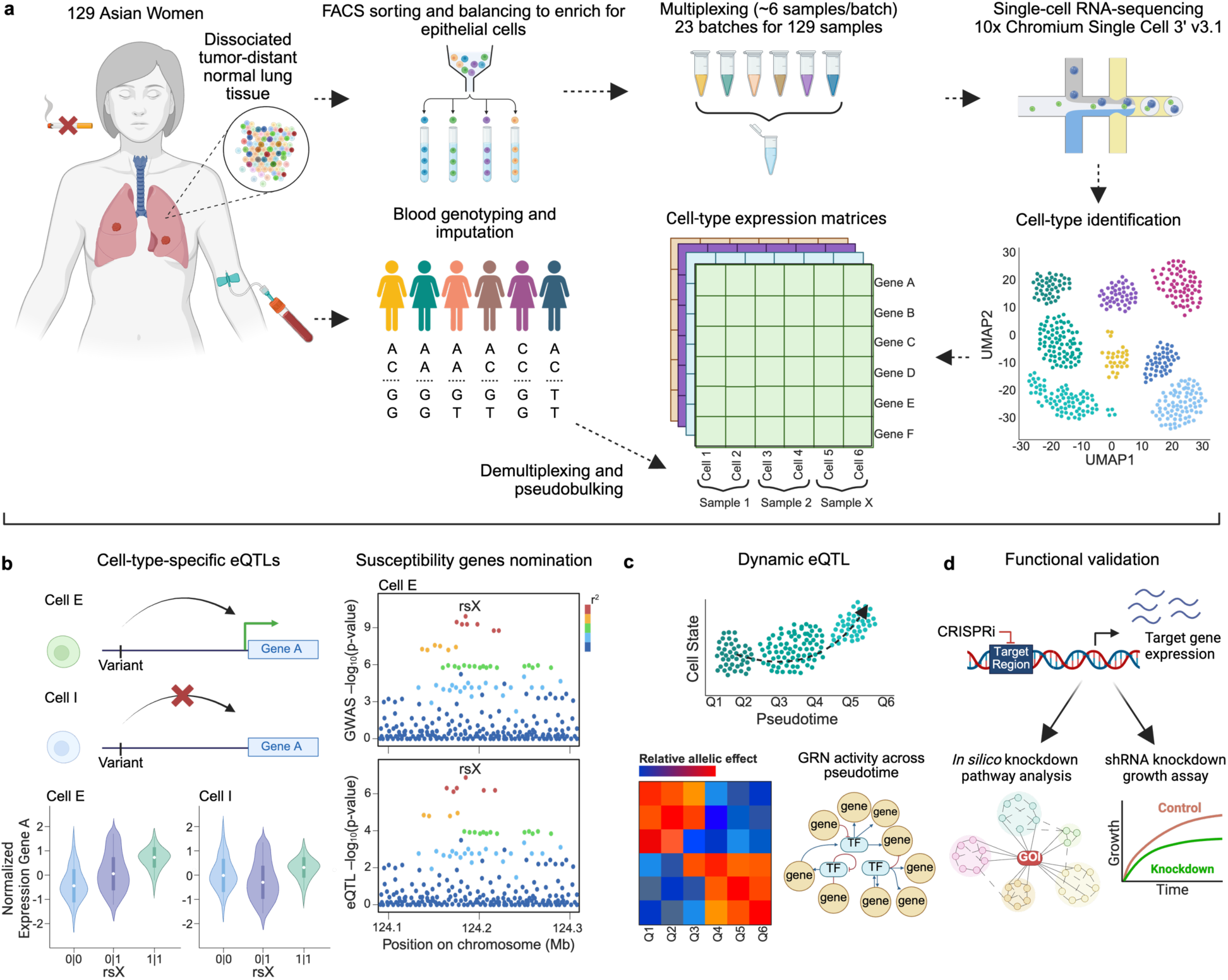
Lung single-cell eQTL dataset of Asian women never-smokers enables elucidation of genetic susceptibility to lung cancer Schematic of study design. **a,** Workflow of establishing expression and genotype data for single-cell lung eQTL analysis, including sample collection, cell type balancing and epithelial cell enrichment, multiplexing and single-cell RNA-sequencing, and DNA genotyping and imputation. **b,** Single-cell eQTL mapping and nomination of context-specific susceptibility genes. **c,** Dynamic eQTL mapping in alveolar epithelial cell states and gene regulatory network (GRN) of dynamic eQTL genes. **d,** Functional validation using CRISPRi targeting candidate causal variants to assess the effect on susceptibility genes, followed by *in silico* pathway analysis and/or growth assay to implicate the roles of these susceptibility genes in lung cancer etiology. FACS: fluorescence-activated cell sort. The figure was created using Biorender.

Following iterative clustering and annotation, we identified 41 cell types (246 - 33,953 cells/cell type) in four main categories: Endothelial, Epithelial, Immune, and Stromal (Figs. 2a-b, Supplementary Fig. 2, Methods, cell type abbreviation listed in Supplementary Table 4).

**Figure 2.**
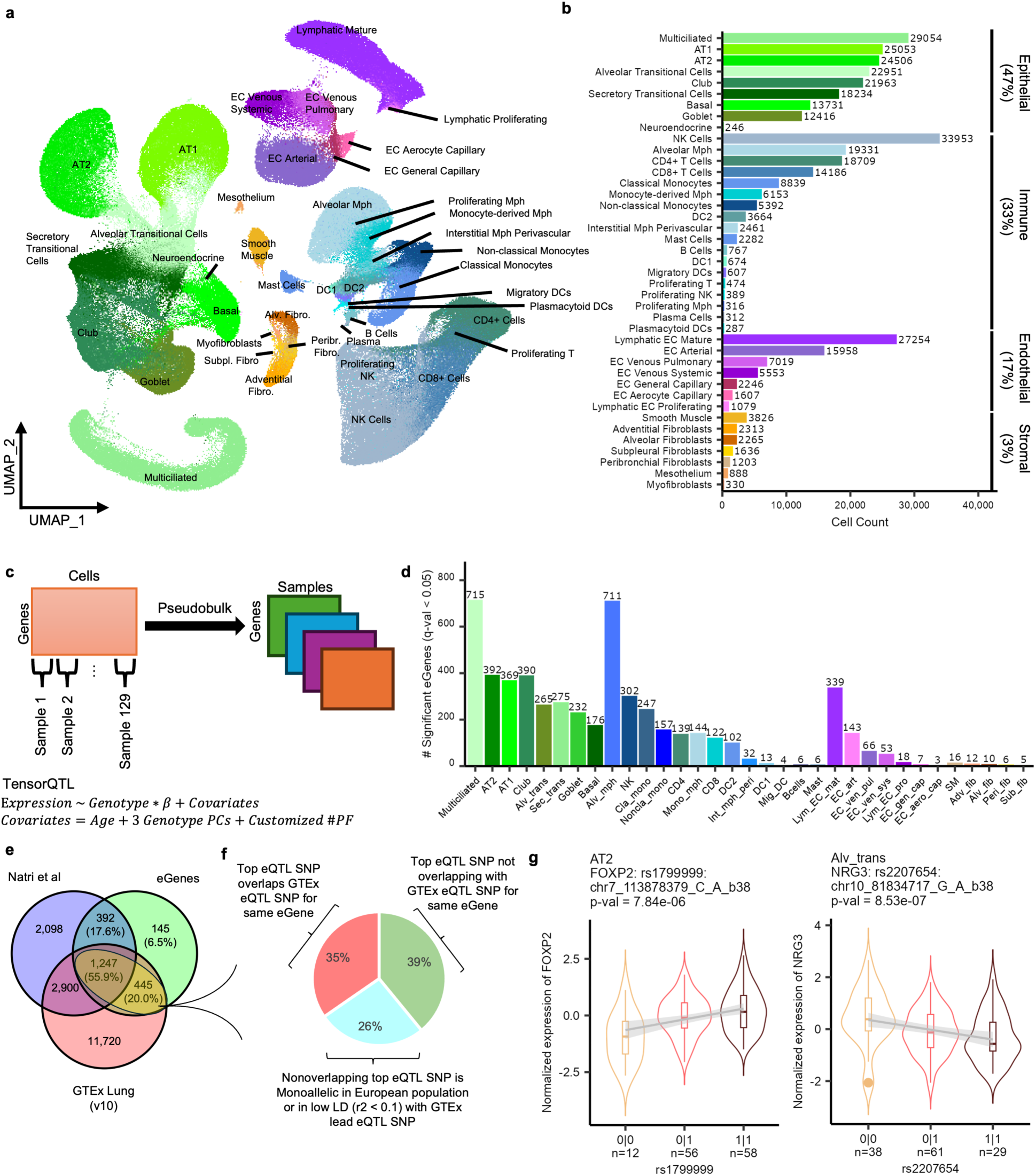
Epithelial cell-enriched lung cell atlas revealed cell type specific eGenes and population specific eQTLs **a,** Uniform manifold approximation and projection (UMAP) plot depicting 41 lung cell types from 129 individuals. **b,** Breakdown of number of cells in each cell type is depicted on the bar plot. Percentage of each cell category is shown on the right. Depiction of each cell category is represented in different shades; green: epithelial, blue: immune, purple: endothelial, golden: stromal. **c,** Single-cell eQTL mapping schematic in 33 cell types. Genes x cells matrices were pseudobulked to genes x samples matrices for each cell type. Covariates for eQTL mapping include age, genotype PCs, and expression PEER factors (PF). **d,** Number of significant eGenes (q-value < 0.05) for each of the 33 cell types are plotted. Color schematic of each cell type and category follows that of **a. e,** Venn diagram showing overlap of our eGenes with two external datasets. Lung GTEX (v10) and recent single-cell eQTL dataset (Natri et al), both external datasets, comprise predominantly European individuals. Percentage of overlap or unique subset is in reference to our dataset. **f,** Pie-chart shows proportion of our top eQTL SNPs (1,692) overlapping or not overlapping with GTEx eQTL SNP for the same eGene. Of non-overlapping, proportion of mono-allelic or low LD is shown. **g,** Violin plots showing association between normalized expression of *FOXP2* and *NRG3* with top eQTL SNP, rs1799999 and rs2207654, in AT2 and Alveolar Transitional cells, respectively. Center line denotes the median, while the 25^th^ and 75^th^ percentile is marked as the lower and upper line of the box, respectively. Whiskers extend 1.5 times from the 25^th^ and 75^th^ percentiles; outliers are represented as dots. The violin width reflects the density. SNP chromosome and position are based on hg38 assembly. 0 = reference allele, while 1 = alternative allele.

Expression of the final cell type marker genes aligned with Human Lung Cell Atlas (HLCA) reference and were independently validated by NSForest^31^ which detects “necessary and sufficient” marker (61% of cell types had >=1 matching marker(s) between HLCA and NSForest) (Supplementary Table 4, Supplementary Fig. 3). Consistent with our design, we observed an enrichment of epithelial cells, which make up approximately 47% of our dataset, followed by immune (33%), endothelial (17%), and stromal (3%) (Fig. 2b). We observed >83% of our final cell types were represented in >87% of individuals, and each single-cell batch, on average, contributed evenly to every initial cluster (∼1/23 batches or ∼4.3%), which suggests minimal batch effect or sampling bias (Supplementary Table 5).

Of note, we detected most of the reported cell types of origin for major lung cancer histological subtypes, including rarer and transitional cell types. Namely, in addition to AT2, AT1, club (LUAD), basal (lung squamous cell carcinoma or LUSC), and neuroendocrine cells (small cell lung cancer or SCLC), we identified a substantial proportion of epithelial cells displaying a transitional nature (∼11% of our dataset). Recent publications have used *SCGB3A2* as a marker for transitional/progenitor cell types in distal airways and alveolae, such as “AT0” and “RAS”. We therefore defined clusters which either moderately or highly expressed this marker to have transition potential^20–22^. We then incorporated other markers to define two transitional cell types, Alveolar Transitional (*SCG3A2+, SFTPC+, SFTPB+*) and Secretory Transitional (*SCG3A2+, SFTPC-, SCGB1A1-*). We confirmed that these transitional epithelial cells are indeed found in our patient samples by performing multi-color fluorescence immunohistochemistry against SFTPC, SCGB3A2, and SCGB1A1 in a subset of matched tissue blocks from our cohort (Supplementary Figs. 4 and 5). In addition to enriching cells of lung cancer origin in the epithelial category, we retained other lung cancer-relevant cell types, such as those in immune cell category^29,32^. Namely, abundant alveolar macrophages as well as distinct cell states and rarer immune cell types were detected, including 4 types of macrophages and 4 types of dendritic cells (Figs. 2a-b). Overall, our dataset efficiently captured relevant contexts of lung tumorigenesis, which enabled us to interrogate the underlying genetic mechanisms.

### Detection of cell type and population specific eQTLs

To investigate how genetic variations influence gene expression regulation in each lung cell type, we performed sc-eQTL analysis. We applied pseudo-bulk eQTL approach using TensorQTL^33^ (Fig. 2c) in each of 33 qualifying cell types, with inclusion criteria similar to those of recent studies^26,34^ (>=40 individuals with 5 or more cells, Supplementary Table 6, Figs. 2a-b, Methods). Among 8,169 qualifying genes (expressed >= 10% cells, >=1 SNPs in +/-1 MB), we detected 2,229 (27.3%) genes with a significant eQTL or eGenes in at least one cell type (q-value < 0.05) (Fig. 2d, median = 122, range = 3 - 715; eGenes/Cell Type, Supplementary Table 7). Permutation analysis by randomly shuffling phenotypes demonstrated that type I errors were tightly controlled (Supplementary Fig. 6). Similar to previous studies^26,28^, the numbers of eGenes positively correlated with the numbers of cells, underscoring the differences in detection power across cell types (Supplementary Fig. 7a). A less pronounced correlation between the number of individuals and detected eGenes was also observed (Supplementary Fig. 7b).

To determine the validity of the detected eGenes (q-value < 0.05), we compared our dataset to eGenes detected in two external datasets, bulk lung eQTL data from GTEx (v10) (601 individuals, ∼85.3% European), and a recent sc-eQTL study^26^ (113 individuals, ∼66.7% European, ∼57.9% with ILD). We found that most (93.5%) of our eGenes were replicated in one of these external datasets (Fig. 2e), whereas 6.5% were unique to our dataset. We observed similar replication rates in the sc-eQTL dataset (73.5%) and the much larger GTEx data (75.9%). This suggests a high similarity between single-cell datasets reflecting cell-type-specific findings even with different statistical approaches (Methods).

While the replication of eGenes validated our findings, we also noted that eGenes unique to our dataset displayed cell type and population specificity. Namely, compared to GTEx-replicated eGenes, our dataset-specific eGenes included a larger proportion of eGenes significant in a single cell type (68% vs 53%). Furthermore, compared to eGenes shared with both mainly European datasets, the top eQTL SNPs associated with our dataset-specific eGenes displayed a higher average MAF in East Asians (1000G v5 EAS MAF 0.41 vs 0.36, Welch’s t-test: p-value = 0.035). For example, *FOXP2* was an AT2-specific eGene unique to our dataset, displaying a higher minor allele frequency (MAF) of top variant, rs1799999 in EAS (0.36) compared to EUR (0.11). Similarly, *NRG3* was an Alveolar Transitional Cell-specific eGene with the top SNP, rs2207654, displaying EAS specificity (MAF EAS: 0.46, EUR: 0.09) (Fig. 2g). These data suggested that lower statistical power due to the low MAF of these SNPs in EUR populations led to failure of detecting these eGenes in mainly EUR eQTL datasets. To glean at potential ancestry specific eQTLs, we compared the eQTL results from common eGenes (1,692) in ours and GTEx results (both using similar statistical approaches for eQTL mapping). We found 1,106/1,692 (65.4%) of our top variants were not detected as an eQTL SNP for the same eGene in GTEx. Of these, 445/1,106 (40.2%) were in low LD (r^2^ < 0.1) with top variant in GTEx or monoallelic in European population (Fig. 2f). These results suggested that our data detected unique signals that were not previously seen in mainly European eQTL datasets. Finally, we compared our eGenes to the recently published Chinese sc-eQTL dataset^28^ and observed that 60% (1,335/2,227) of our eGenes were not detected in Fu et al data, including 83% of our dataset-specific eGenes (relative to the two European datasets). Our unique findings compared to Fu et al could be due to different sample processing such as our epithelial enrichment as well as different cell type annotation (33 vs 17 cell types for eQTL detection) and eQTL mapping approaches. These results highlighted that our dataset could identify unique eGenes, including some that are potentially cell-type-specific as well as East Asian-specific eQTLs that were not detected in previous datasets.

### Cell type and lineage-specific *cis*-regulation of lung

To assess whether the detected eQTL SNPs represent cell-type-specific gene regulation mechanisms, we performed functional enrichment and cell type sharing analyses. First, we asked if general lung-relevant *cis*-regulatory elements are enriched in the dataset-wide eQTL SNPs using ENCODE ChromHMM 15 chromatin state annotation (E096, Lung). “Pseudo-lung” eQTL SNPs (LD-pruned and collapsed from all cell types) were significantly enriched with the largest odds ratio for Enhancers, followed by Flanking Active TSS, and Active TSS compared to non-eQTL SNPs. In contrast, eQTL SNPs were enriched with lower odds ratios or depleted for Quiescent/Low, Heterochromatin, and Weak Repressed PolyComb regions (Fig. 3a). This data suggests that our eQTL SNPs have a role in regulating gene expression with a subset potentially functioning as enhancers.

**Figure 3.**
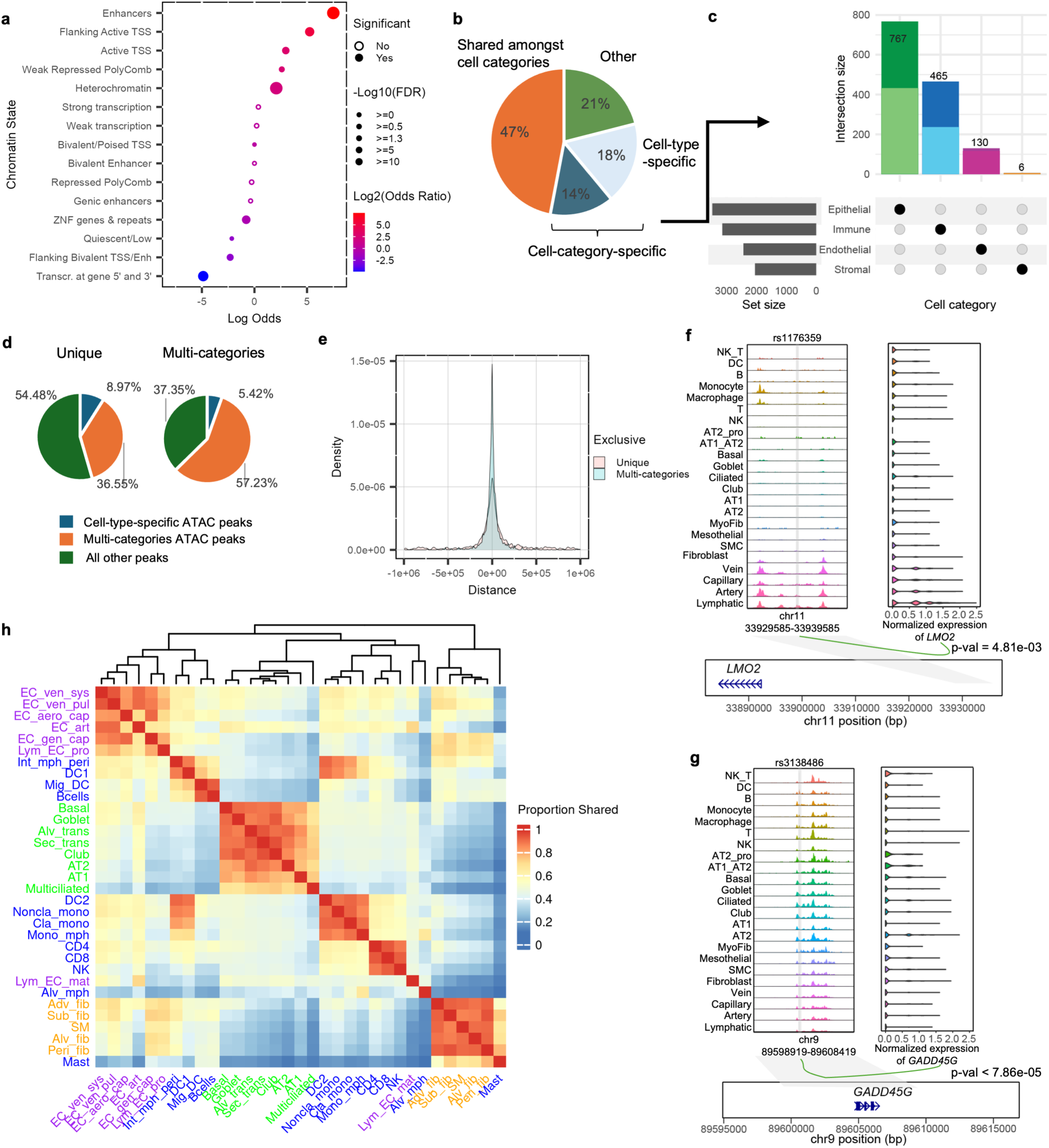
Cell-type-specificity, sharing, and functional implication of eQTL SNPs **a,** Enrichment or depletion of LD-pruned eQTLs combined across the cell types (q-value < 0.05, TensorQTL) in ENCODE ChromHMM 15-state annotation (Lung, E096). Log_2_ (odds ratio) is scaled from blue for depletion to red for enrichment. Size of circle represents log_10_ (FDR), where significant hits as defined by FDR < 0.05 are solid circles, while non-significant are empty circles. **b,** Pie-chart showing shared or cell-type-specificity of top eQTLs (q-value < 0.05, TensorQTL) following mashr correction (significance defined by lfsr < 0.05). **c,** Upset plot showing the number of cell-category-specific top-eQTLs, and a proportion, which is cell-type-specific, is shown in a light shade. **d,** Pie-charts depicting relative percentage of cell type-specific eQTL SNPs overlapping cell-type-specific, shared (across all four cell categories), or remaining ATAC-peaks. **e,** Density plot showing distance from cell-type-specific or shared (across all four cell categories) eQTL SNPs to the TSS of their target genes. **f-g,** Sequencing tracks representing chromatin accessibility of lung cell types, where each track represents the aggregated snATAC signal of each cell type normalized by the top number of reads in the region (0-210 top, 0-370 bottom). Grey vertical lines represent the location of variants of interest. rs1176359, a lymphatic cell-specific eQTL SNP overlaps a lymphatic cell-specific ATAC peak, while shared eQTL SNP, rs3138486, overlaps shared ATAC peaks. Loops show significant linkage between ATAC peak accessibility and gene expression by Signac with their p-values (Long et al). **h,** Heatmap of proportion of effect size sharing between cell types. To be considered shared, the eQTL is significant (lfsr < 0.05) in both tested cell types and effect is in the same direction and within 0.5 to 2-fold.

As enhancers are thought to be cell type specific, while promoters are shared across multiple cell types^35–37^, we aimed to first characterize eQTL sharing and exclusivity between cell types. To alleviate the detection power differences between cell types, we utilized mashr^38^, which incorporates information across conditions (cell types) to refine effect size estimates and improves statistical power when signal is shared across cell types. Of 4,372 unique top eQTLs (top eQTL SNP for an eGene in each cell type) that we considered, 4,274 (97.8%) were significant after mashr correction (local false sign rate or lfsr < 0.05). In terms of significance of these SNPs across cell types, the number of significant eQTLs for smaller cell types increased (Supplementary Fig. 7), while single-cell type eQTLs decreased (Supplementary Fig. 8b). Based on these criteria, we observed that 47% (2,002) were shared among cell types across two or more cell categories. On the other hand, 32% (1,368) were only shared between cell types in the same cell category, while 18% (787) were exclusive to a single cell type (Figs. 3b-c, Supplementary Fig. 8a).

Next, we integrated this cell-type-specific and shared eQTL SNPs with cell type resolved *cis*-regulatory elements using single-cell ATAC-seq data of similarly processed lung tissues from our recent study^29^. We overlapped eQTL SNPs with different ATAC-peak groups (cell-type-specific, multi-categories, and all others) and observed a significantly different overlap proportions across the three ATAC-peak groups between cell-type-specific and shared eQTLs (Fig. 3d, χ^2^ test, p-value = 1.68e-04). We found that eQTL SNPs shared across cell categories were closer to transcription start sites (TSS) of their associated eGenes and mostly overlapped with ATAC-seq peaks common to multiple cell categories (57% of shared eQTL SNPs vs 36% of specific eQTL SNPs). At the same time, cell-type-specific eQTL SNPs were further away from TSS and more frequently overlapped with cell type exclusive peaks (9% of specific eQTL SNPs vs 5% of shared eQTL SNPs) (Figs. 3d-e). This finding is consistent with previous observations that shared ATAC peaks were more likely to contain promoter elements whereas exclusive peaks contain distal enhancers^29,39^. Together, our data suggests that cell-type-specific eQTL SNPs are functioning likely as a part of enhancers, while shared eQTL SNPs likely as a part of promoters. For example, a cell-type-specific eQTL SNP, rs1176359, overlapped a Lymphatic-specific ATAC-seq peak, which is ∼70,000 bp upstream of its associated Lymphatic-specific eGene, *LMO2*, suggesting an enhancer function (Fig. 3f). Conversely, an eQTL SNP shared across all 4 categories, rs3138486, overlapped an all-category-shared peak next to the TSS of *GADD45G,* suggesting promoter function (Fig. 3g). The chromatin accessibilities of these two ATAC-seq peaks were also significantly correlated with the respective eGene expression levels (*LMO2*, p-value = 4.81e-03; *GADD45G*, p-value = 7.88e-05) based on peak-gene linkage from our previous study^29^.

Since regulatory mechanisms can be shared between cell types of the same cell lineage^18^, we further assessed pair-wise effect size sharing patterns across lung cell types. We observed that effect size sharing of allelic direction and magnitude was predominantly observed between cell type pairs within a cell category and sub-category lineage. Among endothelial, epithelial, and stromal categories, the median sharing within category was 37%. Intriguingly, the immune category had very little sharing, which may speak to the different regulatory mechanisms in its diverse cell types. However, within immune sub-categories, there was a higher level of sharing. For example, among lymphoid cell types (NK, CD4 and CD8 T cell), effect size sharing was 74.1% (Fig. 3h). Altogether, our identified cell-type-specific and shared eQTLs capture genetic regulation underlying distinct lung cell categories and represent functionally relevant *cis*-regulatory mechanisms.

### GWAS integration highlights the role of alveolar cells in lung cancer etiology

Having established the cell type relevant roles of lung sc-eQTLs, we integrated them with lung cancer GWAS data to identify cell type relevant susceptibility genes. For this, we first performed colocalization analyses using summary statistics from two recent lung cancer GWAS from diverse populations^13,14^ (Methods). In total, we identified 12 colocalized genes from 10 loci in 11 cell types (coloc^40^ PP.H_4_ ≥ 0.70; posterior probability that the two traits share the same causal variant^44^) (Fig. 4a, Supplementary Table 8). These included 4 genes (*ACTR2, ROS1, TCF7L2*, and *CAT*) that have not been identified from published coloc or TWAS results using the same GWAS statistics with bulk lung eQTL data^13,17,41^.

**Figure 4.**
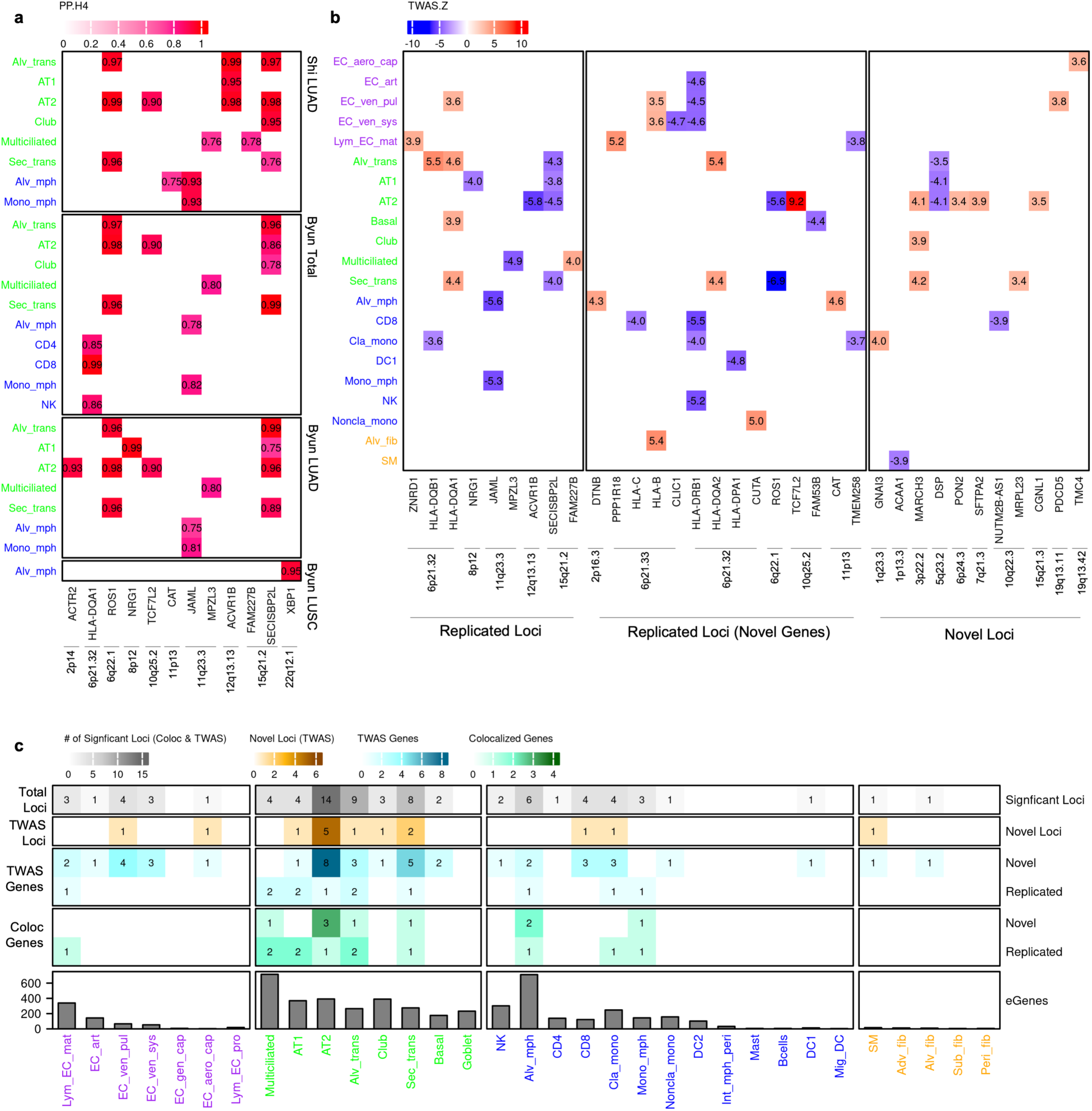
GWAS integration highlights the role of alveolar cells in lung cancer etiology a, Heatmap of colocalized genes (defined by PP.H4 > 0.70) from 4 GWAS summary statistics with lung cancer subtype risk indicated. Shi et al is from EAS population, whereas Byun is multi-ancestry. Epithelial cells are in green, while immune cells are in blue. **b,** TWAS analysis using Shi et al. (LUAD) summary statistics. Heatmap shows TWAS z-scores of significant genes (FDR < 0.05), whereby blue indicates low expression is associated with risk, while red indicates high expression is risk-associated. Results are divided into replicated loci and genes, replicated loci with new genes compared to previous coloc and TWAS using bulk tissues, and novel TWAS-identified loci. Endothelial cells are in purple, epithelial in green, immune in blue, and stromal in golden. **c,** Summary of coloc and TWAS analysis. Cells in their respective categories are ordered by number of cells (high to low), with number of eGenes shown. The number of replicated and novel coloc genes are shown in green, while TWAS genes are in blue. Number of novel loci as defined by TWAS is in brown, and the number of total significant loci from coloc and TWAS combined are in grey.

**Figure 5.**
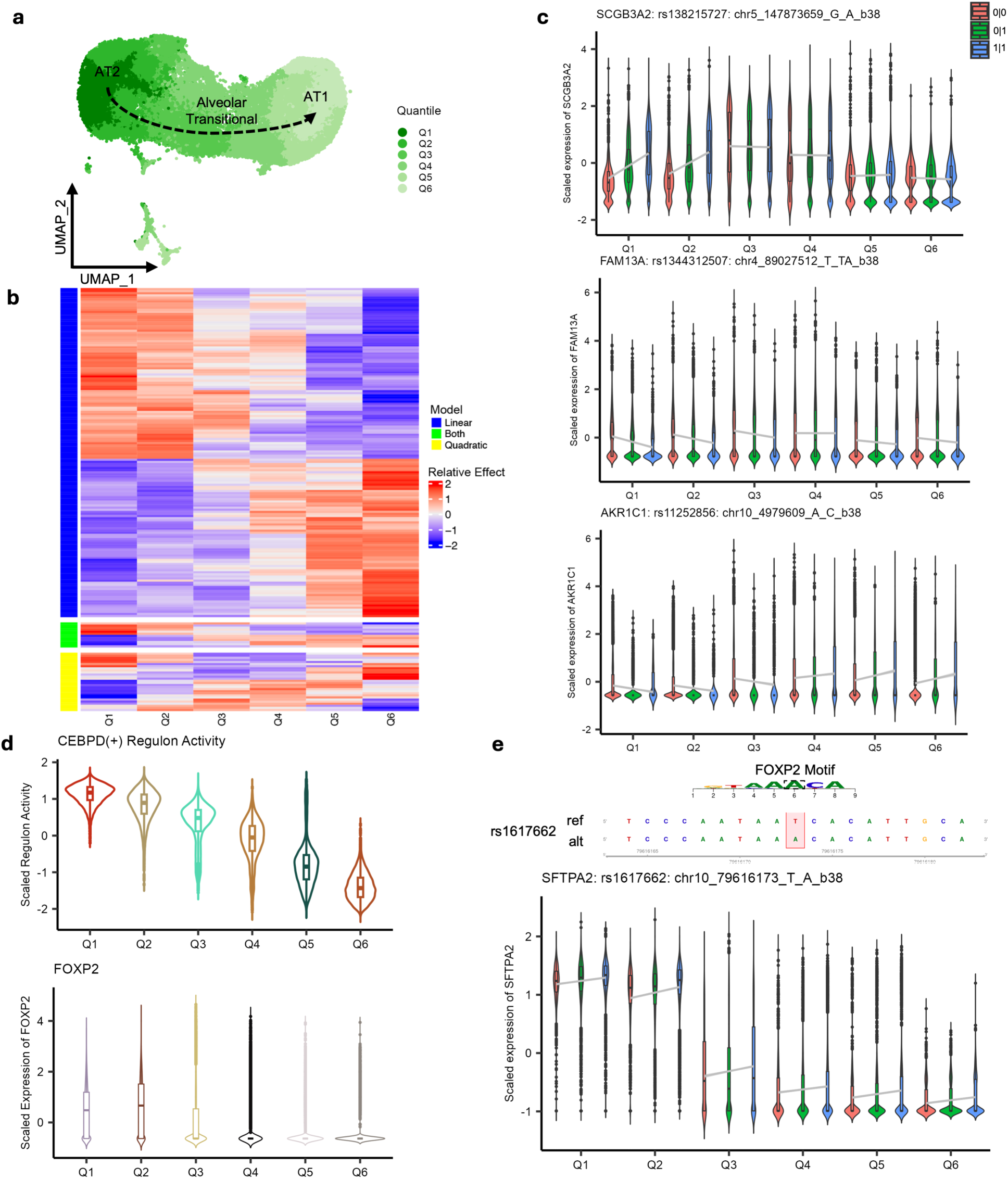
Trajectory analysis reveals dynamic gene programs in alveolar epithelial cells a, Monocle3 UMAP plot of epithelial cell types; 6 quantiles are used to represent the cell states from AT2 (∼Q1, Q2) to Alveolar Transitional (∼Q3, Q4) to AT1 (∼Q5, Q6). **b,** Heatmap of relative allelic effect size of significant dynamic eQTLs which fit linear, quadratic, or both models. **c-e**, Violin plots, where the center line denotes the median, while the 25^th^ and 75^th^ percentile is marked as the lower and upper line of the box, respectively. Whiskers extend 1.5 times from the 25^th^ and 75^th^ percentiles, outliers are represented as dots. The violin width reflects the density. **c,** Association of normalized and scaled gene expression with their significant interaction eQTL SNPs through the cellular states (6 quantiles). Three representative genes fitting a linear (*SCGB3A2, FAM13A, AKR1C1*) model of allelic effect change are shown. Chromosome and position of the variants are based on hg38. 0 = reference allele, while 1 = alternative allele. **d,** Top plot is scaled Regulon activity of *CEBPD(+)* through cell states. Bottom plot shows normalized expression of *FOXP2*, which is a part of the *CEBPD(+)* regulon. **e,** Association of normalized expression of *SFTPA2* with interaction eQTL, rs1617662, across the cell states, rs1617662 is part of a binding motif for FOXP2, where the alternative allele A has the consensus sequence, while the reference allele, T, is predicted to have a weaker binding, based on motifbreakR.

To improve detection power beyond the known GWAS loci, we performed transcriptome-wide association study (TWAS) using FUSION^42^ by aggregating the effects of multiple variants for the ancestry-matched LUAD GWAS data^14^ (Methods). TWAS nominated 34 susceptibility genes across 20 loci in 21 cell types (Fig. 4b, Supplementary Table 9). While nine of these genes were a replication of previously published coloc or TWAS genes using bulk lung eQTL data in any population, 25 (73.5%) were genes that have not been reported from known GWAS loci or from the loci that did not reach genome-wide significance in the GWAS (10 loci). Of the 34 genes, 10 were replicated from our colocalization analysis.

TWAS and colocalization identified susceptibility genes from every cell category, with the majority in epithelial and immune cells. Of the 11 cell types with a colocalization, 6 were of epithelial origin while 5 were immune, underscoring the roles of epithelial and immune cells in lung cancer etiology^29^. Similar to colocalization, half of our TWAS genes were found in epithelial cell types. Notably, *ROS1* (6q22.1) and *TCF7L2* (10q25.2) displayed the lowest and highest Z-scores, respectively (Fig. 4b), and were consistently detected by both coloc and TWAS in AT2 and/or transitional cells. Although *ROS1* fusion is a known somatic driver of LUAD, especially in never-smoking women^43^, physiological function of *ROS1* remains underexplored^44^. Notably, previous functional studies of the 6q22.1 locus or bulk lung tissue eQTL data have focused on a neighboring gene, *DCBLD1*^17,45^, with lack of direct evidence for *ROS1*. *TCF7L2* in 10q25.2 locus, which is an effector of the Wnt/β-catenin pathway was identified as a susceptibility gene exclusively in AT2 cells. This gene is an eGene only in AT2, although it is highly expressed in multiple other cell types with high cell numbers, including Alveolar macrophages and AT1 (Supplementary Fig. 9a). Notably, MAF of the top eQTL SNP for *TCF7L2*, rs11196100, is 0.26 in East Asian population but lower (MAF = 0.05) in European population, thereby limiting statistical power for eQTL detection in European datasets. Consistent with this observation, *TCF7L2* is not an eGene in GTEx lung (v10) tissue but an eGene in AT2 cells in Natri et al^26^ where the lead SNP is in low LD with our lead SNP (r^2^ < 0.01 in EAS). For immune cells, findings from both colocalization and TWAS were related to stress and immune response (e.g. *HLA* genes, *CAT*, and *JAML*). TWAS also identified susceptibility genes from endothelial and stromal cell types. The majority of the findings in endothelial and stromal cells were from the HLA locus; however, we also observed significant genes with a cell-type relevant function, such as *CLIC1*, encoding chloride channel protein, and *ACAA1*, encoding a metabolic protein, in Endothelial venous systemic and Smooth muscle cells, respectively. *CLIC1* has been shown to regulate endothelial cell migration and proliferation and mediate blood vessel formation^46^.

*ACAA1* encodes an enzyme involved in fatty-acid beta-oxidation, which can generate acetyl-CoA for energy production.

A summary of our colocalization and TWAS analyses is shown in Fig. 4c (Supplementary Table 10). Of the novel susceptibility genes, 50% (13/26) were from GWAS loci identified in Shi et al^14^ (EAS, LUAD), highlighting the utility of ancestry-matched TWAS analysis. Intriguingly, although they are not the most abundant cell types of their respective cell categories; AT2, Alveolar transitional, and Alveolar macrophages had the highest numbers of significant loci, explaining 68% (15/22) of our coloc and TWAS loci. These data underscore the contributions of the cells of the alveoli in lung cancer risk, particularly lung adenocarcinoma.

### Dynamic genetic regulation in alveolar epithelial cell states relevant to lung cancer

The GWAS integration highlighted the contributions of AT2 and Alveolar transitional cells in lung cancer etiology, particularly in LUAD (Fig. 4c). This is consistent with the transitional nature of AT2 cells as a facultative stem/progenitor cell, replenishing AT2 and AT1 populations upon injury, and a cell type of origin for LUAD^23,47^.

To glean whether genetic regulation associated with these changing alveolar cellular states contributes to lung cancer risk, we performed dynamic eQTL analyses. We first estimated alveolar epithelial cell state trajectory from AT2 (pseudotime = 0) via Alveolar Transitional to AT1 (pseudotime ≈ 30) (Monocle3^48,49^, Methods) (Fig. 5a) and further evenly partitioned them into 6 quantiles, a similar approach to previous studies^24,50^. We then performed interacting eQTL analyses for the top 4,372 eQTLs across the 6 cell states using negative binomial regression (Methods). Among these, 785 eQTLs (494 genes) showed significant interaction (int-eQTL, FDR < 0.05), indicating dynamic allelic effect across alveolar epithelial cell states (Supplementary Table 11). Ten of these 494 genes overlapped the genes identified from colocalization or TWAS in this study, suggesting that 28% of lung cancer susceptibility genes could be affected by alveolar cell state contexts.

Next, we asked whether some of these 785 int-eQTLs reflect cell-state-regulating gene expression programs. For this, we first identified 198 int-eQTLs significantly fitting either a linear (79.8%), quadratic (14.1%), or both models (6.1%) (p-value < 0.05) (Fig. 5b) and focused our analysis on 103 genes associated with the linear int-eQTLs, which were the most prevalent.

Among those fitting a linear model, we found *SCGB3A2*, a marker for transitional epithelial cells^20–22^, showing linear change of allelic effect for rs138215727; notably, the significant allelic effect in AT2 states disappears in alveolar transitional cells, although the levels are still high in the transitional stage (Fig. 5c). Similarly, we observed a strong allelic effect for *FAM13A* in AT2 cell state, which flattens out in later stages. Notably, this gene is historically linked to COPD risk by mediating signal transduction, but recently was associated with lung cancer risk^28,51–53^. On the other hand, for *AKR1C1*, a gene encoding aldo-keto reductase and implicated in metabolic reprogramming and cancer cell proliferation, allelic effect direction flips from AT2 to AT1 cell states (Fig. 5c)^54^.

To nominate potential *trans*-regulators mediating these linear int-eQTL effects, we further detected transcription factor (TF)-centered gene regulatory networks (regulons) in the three alveolar cell types using *SCENIC*^55^ (Methods). From this we detected 119 regulons displaying a significant linear activity across the six cell states. Among them, 87 genes associated with 62 regulons overlapped the genes with an int-eQTL effect fitting a linear model (Supplementary Fig. 10a). Pathway analysis of these 87 genes and 62 TFs highlighted lung epithelial development, Wnt signaling, and integrated stress response, suggesting that this subset reflects potential cell-state-regulating genes and associated int-eQTL genes (Supplementary Fig. 10b).

Among these 87 genes were 4 lung cancer susceptibility genes, including *SFTPA2*, which was identified as a TWAS gene in AT2 cells from a locus not achieving genome-wide significance in GWAS. To identify potential cell-state-associated TFs mediating *SFTPA2* int-eQTL, we explored TF binding motifs affected by 32 proxy SNPs of the interacting SNP, rs1240314687 (r^2^ > 0.8). Among 13 motif-altering SNPs identified by motifbreakR^56^ (p-value < 1e-04), we observed that rs1617662 alters a binding motif of FOXP2. Notably, *FOXP2* alongside *SFTPA2* were part of the *CEBPD* regulon, both displaying an expression correlation with *CEBPD* regulon activity, nominating FOXP2 as a candidate mediator of *SFTPA2* int-eQTL (Figs. 5d-e). Moreover, the effect size change of rs1617662 on *SFTPA2* expression resembles *FOXP2* expression change across the cell states. Overall, our data revealed that gene regulatory programs that are dynamically changing through the alveolar epithelial cell states might affect dynamic eQTLs, including those relevant to lung cancer susceptibility.

### *TCF7L2* is an AT2-specific susceptibility gene in East Asian LUAD locus at 10q25.2

To experimentally validate the LUAD susceptibility genes identified in alveolar epithelial cells, we focused on *TCF7L2* and *ROS1*, two genes with both TWAS and colocalization support in AT2 cells and from the loci originally identified in Asian never-smoking female GWAS^57^ (Figs. 4a-b).

For *TCF7L2* eQTL in AT2, colocalization was observed for rs11196089 (chr10:112,749,531), top SNP in the 10q25.2 LUAD GWAS locus (East Asian)^14^, which also showed a higher MAF in EAS than EUR (0.30 vs 0.05) (Figs 6a-b). Here, the risk-associated C allele is correlated with higher expression of *TCF7L2* (Fig. 6c). To nominate candidate causal variants (CCVs) of LUAD from this locus, we selected 42 SNPs with log likelihood ratio (LLR) < 1:1,000 in GWAS statistics or R^2^ > 0.8 in reference to r11196089 in East Asian populations.

**Figure 6.**
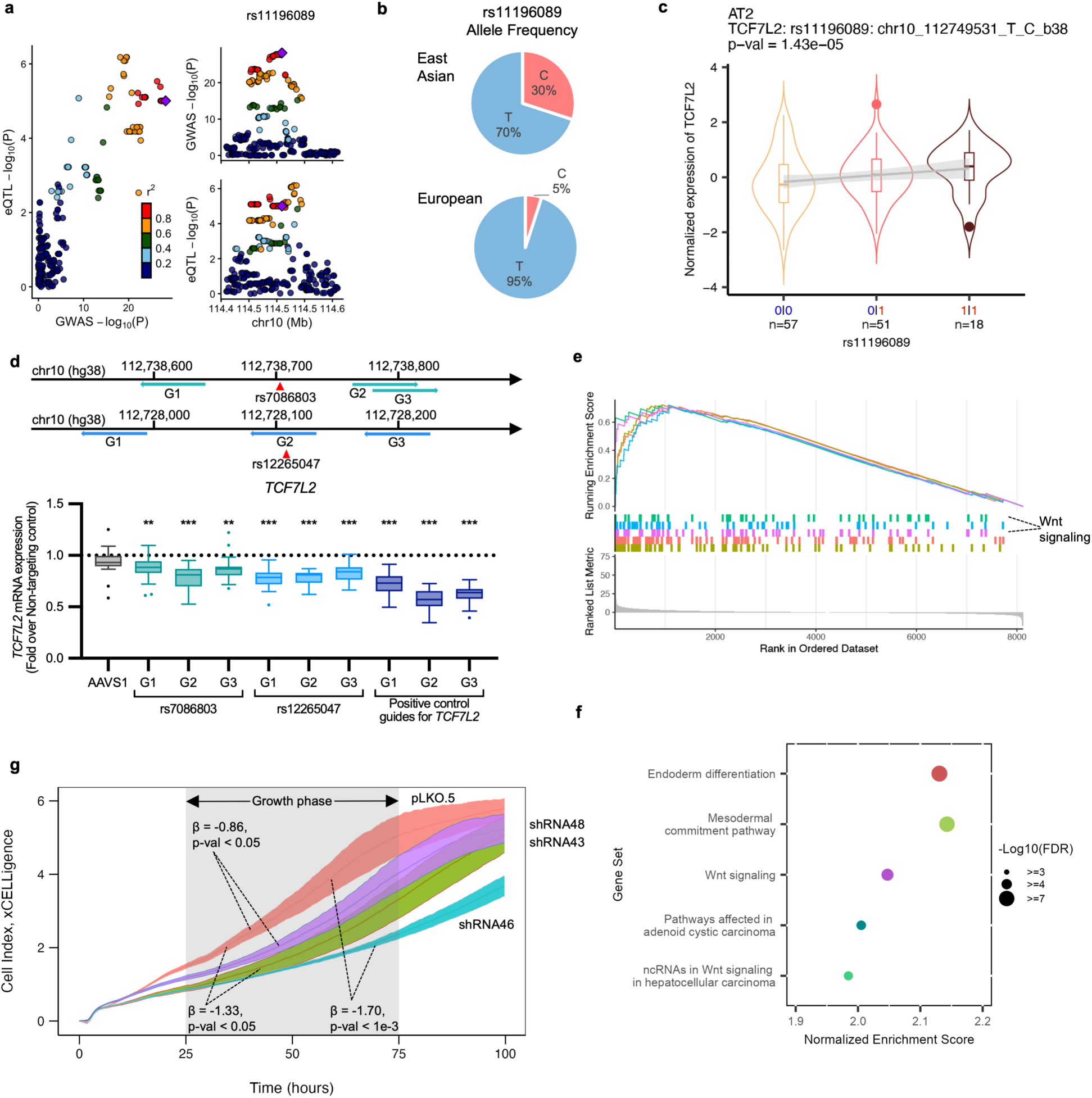
*TCF7L2* is a LUAD susceptibility gene involved in Wnt signaling and promoting cellular growth in the AT2 lineage **a,** Locus zoom plots of eQTL in AT2 cells for *TCF7L2* and GWAS (Shi et al EAS, LUAD) p-values of the region (+/-100 kb) around the GWAS top SNP, rs11169089 in the 10q25.2 locus. LD (r^2^) relationships between the variants are extracted from 1000G EAS Phase 3 v5. **b,** Allele frequencies difference between East Asian and European from 1000G **c,** Association between normalized expression of *TCF7L2* and rs1196089 is shown as violin plots. 0 = reference allele, while 1 = alternative; red indicates risk allele, while blue indicates protective. Center line denotes the median, while the 25^th^ and 75^th^ percentile is marked as the lower and upper line of the box, respectively. Whiskers extend 1.5 times from the 25^th^ and 75^th^ percentiles; outliers are represented as dots. The violin width reflects the density. Note the nominal p-value threshold is 1.31e-05. **d,** Location of guide RNAs targeting the regions around our SNP(s) of interest. Tukey plot shows *GAPDH*-normalized mRNA levels of *TCF7L2* in A549 cell line from 6 replicates from three independent experiments (n =18). Fold change of target gene expression over non-targeting control is shown as median with interquartile range (IQR) in a box. Whiskers extend 1.5 times IQR, with outliers shown as dots. AAVS1 represents a safe harbor site-targeting gRNA. P-values were calculated using a two-tailed Mann Whitney U test. **e** and **f,** GSEA analysis in AT2 cells comparing individuals that highly (75^th^ percentile) vs lowly (25^th^ percentile) expressed *TCF7L2*. The top 5 most enriched gene sets by normalized enrichment score (NES) are shown. A positive NES indicates enrichment of the gene set compared to the reference. Cumulative enrichment score (e) and NES (f) for the top 5 gene sets are shown with a matching color for each gene set in GSEA (e) and NES (f) plots, where circle sizes are scaled to the-log_10_ FDR values. **g,** A representative xCELLigence growth assay result among 5 independent experiments, where mean cell index with SEM of up to 4 replicates over time is shown. Beta estimates (during growth phase) from linear-mixed-effect models comparing shRNA(s) to control are shown; negative sign is indicative of decreased growth rate compared to control. *** denotes p-value < 1e-04 while ** p-value < 1e-03.

These variants were tested for allelic transcriptional activity in two lung cancer cell lines as part of massively parallel reporter assays (MPRA)^58^, and 14 of them displayed significant allelic difference (FDR < 0.01) in one or both lung cancer cell lines. Of these 14, only rs7086803 and rs12265047 were located within an enhancer histone mark (ENCODE ChromHMM, lung tissues or cell lines, Supplementary Fig. 11). Further eQTL fine-mapping using *susie*^59^ (Methods) demonstrated that both rs7086803 and rs12265047 were in eQTL credible sets, nominating them as likely functional variants contributing to both GWAS and eQTL signals. To test the functional effect of these SNPs on *TCF7L2* expression, we performed CRISPR interference (CRISPRi) targeting the regions on or near these SNPs, using three guide RNAs (gRNAs) for each SNP, in A549 (LUAD cell line). We observed that targeting rs7086803 or rs12265047 consistently resulted in 16% (range: 13-22%) or 20% (range: 17-22%) mean reduction of *TCF7L2* expression, respectively, which was slightly lower than the gRNAs targeting the promoter region of *TCF7L2* (Fig. 6d, 37% mean reduction with a range of 28-43%) (p-value < 1e-03 for all guide RNAs except AAVS1 negative control P = 0.054, n = 18, 3 independent experiments combined). These results demonstrated functional relationships between rs7086803/rs12265047 and *TCF7L2* (Fig. 6d).

We then performed *in silico* overexpression and pathway analyses, which indicated that genes co-expressed with *TCF7L2* in AT2 cells are enriched for Wnt signaling and developmental/differentiation pathways (Figs. 6e-f, Methods). The data is consistent with the role of TCF7L2 as an effector of the Wnt/β-catenin pathway. Moreover, Wnt signaling pathway was enriched among the alveolar cell-state-related regulons peaking at AT2 cell states, including int-eQTL genes (e.g., *CAPN8, FRMD4B, MAFF*, and *MAN2A1*) and associated regulons (e.g., FOXP2, NFKB1, and TCF7L1) (Supplementary Fig. 10b). In line with the reported function of *TCF7L2* in cell cycle progression and subsequent growth of colorectal cancer cells^60^, our data suggested that TCF7L2 may have a role in driving proliferation of lung alveolar epithelial cells. To test whether *TCF7L2* impacts lung adenocarcinoma cell growth, we utilized shRNAs targeting three separate regions of *TCF7L2* in A549 cells and subsequently measured cell growth using xCELLigence. We observed that all three shRNAs resulted in decreased levels of TCF7L2 (Supplementary Fig 9b). Consistently, targeting *TCF7L2* resulted in significantly reduced cellular growth compared to control for all three shRNAs in 3-5 independent experiments (median beta across 5 sets for three shRNAs are-0.86,-1.52,-2.38; Fig 6g, Supplementary Table 12). This result suggested that high levels of *TCF7L2* may promote cellular growth in lung adenocarcinoma cells.

We took a similar approach for *ROS1*, whose expression is strongly associated with LUAD risk in alveolar and transitional epithelial cells. Colocalization was observed between *ROS1* eQTL in AT2 and rs6937083 (chr6:117,464,145), top SNP in the 6q22.1 LUAD GWAS locus (East Asian)^14^, where the risk-associated A allele is correlated with lower expression of *ROS1* (Supplementary Figs 12a-b). We nominated 13 CCVs using the same criteria as above, where only 3 displayed significant MPRA allelic difference (FDR < 0.01) only in A549^58^. Of these 3, only rs5879422 showed the same allelic direction as *ROS*1 eQTL and was also located within an enhancer histone mark (Encode ChromHMM, lung cell lines), nominating it the most likely functional variant (Supplementary Figs. 13a-b). To test the functional effect of this SNP on *ROS1* expression, we performed CRISPRi targeting the region on or near this SNP in a LUAD cell line with a moderate *ROS1* expression (H1975; DepMap (https://depmap.org/portal/)). We observed that targeting this SNP consistently resulted in 24% (range: 20-29%) mean reduction of *ROS1* expression on a par with the gRNAs targeting the promoter region of *ROS1* (23% mean reduction with a range of 15-34%) or another CCV within the same enhancer element with no allelic effect in MPRA (27% mean reduction with a range of 24-32%, Supplementary Fig. 13c) (p-value < 1e-04 for all guide RNAs except AAVS1 negative control P = 0.02, n = 18, 3 independent experiments combined). Our results demonstrated a functional relationship between rs5879422 and *ROS1* (Supplementary Fig. 12c). To glean how lower expression of *ROS1* may contribute to LUAD risk, we performed a similar *in silico* knockdown analysis.

Intriguingly, we observed that lower expression of *ROS1* is associated with enrichment of oxidative phosphorylation (OXPHOS) pathways in AT2 cells (Supplementary Fig. 12d). This implies that *ROS1* may be involved in OXPHOS pathways, and that altered *ROS1* expression may dysregulate metabolic function which is one of the hallmarks of cancer^61^, suggesting a potential mechanism how *ROS1* can contribute to LUAD risk.

Collectively, using a combination of colocalization, TWAS, functional-annotation, and fine-mapping combined with CRISPRi and MPRA approaches, we nominated CCVs and validated functional relationships between them and their target susceptibility genes. Our *in-silico* and experimental investigations provide clues to how *ROS1* and *TCF7L2* may contribute to LUAD risk.

## Discussion

In this study, we performed eQTL mapping and identified *cis*-eQTLs in 2,229 eGenes across 33 lung cell types from an Asian women never-smoker cohort. Our eQTLs were robust with a high replication rate in external datasets at the gene level, whereas dataset-specific findings highlighted genetic signals more pronounced in Asian populations. Moreover, by incorporating a companion single-cell ATAC-seq dataset^29^, we showed that cell-type-specific eQTL SNPs aligned with matched functional elements, highlighting cell type and lineage-specific genetic regulation. By integrating two recent GWAS summary statistics, we nominated 36 lung cancer susceptibility genes with a striking concentration in alveolar epithelial cells. This finding led to the identification of eQTLs interacting with transitioning alveolar cell states involved in lung adenocarcinoma origin. Unique strengths of our dataset were further highlighted in two experimentally validated LUAD susceptibility genes from East Asian GWAS loci that were not evident when using European or bulk lung tissue-based eQTLs.

Intriguingly, alveolar cells contributed the most to lung cancer risk loci, especially cells of LUAD origin and alveolar macrophages implicated in LUAD tumorigenesis. This finding is consistent with the fact that a larger proportion of our susceptibility gene discovery is based on the ancestry-matched TWAS analysis of LUAD, the most prevalent subtype, especially in East Asia. Although we enriched epithelial cells to compensate for the selective loss during sample processing, detected susceptibility genes could not simply be correlated with statistical power reflected in cell numbers or eGene numbers. For example, AT2 and Alveolar macrophage had the most nominated susceptibility genes in their respective cell category, even though they were not the most abundant cell types in their categories, furthermore AT2 cells had the most significant loci even though it does not have the highest number of detected eGenes (Fig. 4c).

Underscoring the power of our study design, even with a moderate sample size, we identified mostly unreported susceptibility genes (72% of identified genes) from lung cancer GWAS loci. Notably, most LUAD TWAS loci with previously unreported genes were identified from GWAS loci that were identified in East Asian populations^14^ (Supplementary Table 10).

These included the loci originally identified in lung cancer GWAS of never-smoking Asian women^63^ (*ROS1* in 6q22.1 and *TCF7L2* and *FAM53B* in 10q25.2) and those from East Asian LUAD GWAS (*DTNB* in 2p16.3) or TWAS (*CAT* in 11p13) studies^14^ with a substantial proportion of never-smoking women. These data suggested that QTL datasets designed to closely represent ancestry and exposure of the study population, and cell type characteristics of the disease could be effective in susceptibility gene identification.

Aside from nominating previously unreported susceptibility genes, our data provided cellular context to known lung cancer susceptibility genes. Consistent with our previous data using cell-type-specific accessible chromatin regions of lung, eQTL-based lung cancer susceptibility genes in this study were mainly from epithelial and immune cell types. Among them, we validated our previous observation in the 11q23.3 locus, showing an interplay of *JAML* and *MPZL3* in immune and epithelial cells, respectively, while providing more fine-grained cell type information: *JAML* is significantly associated in macrophages (alveolar and monocyte-derived) while *MPZL3* is significantly associated in multiciliated cells. In the 8p12 locus, *NRG1* was previously identified using bulk lung eQTL and later implicated in epithelial cell-specific regulatory connection^29^. Here we identified significant genetic regulation only in AT1 cells, one of the proposed cell types of LUAD origin^62^.

From our TWAS and int-eQTL analyses (Figs. 4b and 5b), we find biological processes underlying both lung cancer and COPD, a well-known lung cancer comorbidity, in alveolar epithelial cells. For example, *DSP* (TWAS) and *FAM13A* (int-eQTL gene), have been traditionally linked to COPD and lung function, but were also recently implicated in lung cancer^28,52,52,63–65^. *DSP* functions in cell adhesion providing structural integrity, whereas *FAM13A* has been shown to regulate different signaling pathways such as β-catenin. Alteration of cell-cell adhesions and β-catenin signal transduction can play a role in both lung cancer as well as COPD and lung function, which is consistent with previous biology, highlighting the utility of our dataset for a broader usage in lung diseases.

The combination of studying an Asian cohort and epithelial enrichment led to intriguing findings in several LUAD loci. The strongest TWAS signals in alveolar cells pointed to two previously underappreciated genes that we successfully validated, *ROS1* and *TCF7L2*. *ROS1* in the 6q22.1 locus was identified in alveolar cell types in our dataset, which may have been missed in a bulk setting without a sufficient proportion of alveolar cells, as this locus was previously associated with *DCBLD1* using bulk lung tissue eQTL^45^. Consistent with our findings, a recent lung single-cell eQTL in a Chinese population also identified *ROS1* in this locus^28^.

Moreover, high-penetrance germline mutations were found in *ROS1* within the known 6q23-25 linkage region from African American families including never-smoking women^66^. Although the role of somatic *ROS1*-fusion in driving lung tumorigenesis is well established, its physiological function and how it may mediate LUAD risk in normal tissues remain enigmatic. Intriguingly, in GTEx (v10) *ROS1* is predominantly expressed in the lung, and in our dataset, *ROS1* is mainly detected in AT2, Alveolar Transitional, and Secretory Transitional (Supplementary Fig. 14). The specific expression in transitional/stem-like cell states suggest that *ROS1* may play a role in self-renewal and differentiation processes. Receptor tyrosine kinases (RTK) respond to paracrine, endocrine, and autocrine signaling to modulate various pathways including metabolic ones^67^. The metabolic needs of a stem-like or transitional cells fluctuate depending on the cell state i.e., quiescent, differentiating, or proliferating. From our *in-silico* knockdown, we observed that lower expression of *ROS1* is associated with enrichment of OXPHOS pathways (Supplementary Fig. 12d). This suggests that *ROS1* may negatively regulate genes that are involved in cellular metabolism, and that lower expression of *ROS1* may promote energy production that could fuel cellular expansion. This conforms to the idea that cells undergoing cellular transformation have increased ATP needs to support rapid cellular proliferation.

Although intriguing, the potential link between *ROS1* and OXPHOS and how it may confer LUAD risk needs experimental validation.

In contrast to *ROS1*, we observed that higher expression of *TCF7L2* is associated with LUAD risk, which is in line with its known function as an effector of the Wnt/β-catenin pathway and promoting proliferation (Figs. 6e-f). Importantly, *TCF7L2* has been previously implicated to the 8q24 multi-cancer locus, where a well-established functional variant, rs6983267, can mediate its allelic binding and in turn affect Myc expression mediating cancer risk^68–71^.

Furthermore, *TCF7L2* is one of the most frequently mutated genes in colorectal cancer, however it has not been directly linked to lung cancer susceptibility. Intriguingly, although this locus, 10q25.2, was initially identified in Asian female never-smoker GWAS, there was no apparent target gene based on bulk tissues QTLs mainly based on European populations, with the nearest gene being *VTI1A*^57^. The lower MAF of the top eQTL SNP in European populations (MAF = 0.05) may have led to its exclusion from the analysis or limited statistical power for eQTL detection. Moreover, *TCF7L2* eQTL was specifically found in AT2 cells of the lung, highlighting alveolar cell-and East Asian-specific signals detected in our dataset. Importantly, we observed that depletion of *TCF7L2* impaired cell growth in A549 (Lung adenocarcinoma cell line, Fig. 6g), suggesting that risk-associated high levels of *TCF7L2* may promote lung cell proliferation. Moreover, in our *in silico* analysis, we observed that high levels of *TCF7L2* in AT2 cells is enriched for the Wnt/β-catenin pathway; this pathway was highlighted in our int-eQTL results (Supplementary Fig.10) to be dynamically changing through alveolar cell states with the peak activity in AT2 states. Altogether this suggests that elevated levels *TCF7L2* can potentially drive proliferation and differentiation of alveolar epithelial cells acting through the Wnt/β-catenin pathway.

Our data suggests that dynamic processes in alveolar cell transition might contribute to lung cancer susceptibility, wherein 10 of our 36 lung cancer susceptibility genes were identified as an int-eQTL gene(s). AT2 cells are thought to be a progenitor cell type in the alveoli, renewing and replenishing alveolar epithelial cells after injury. Given its stem-like nature, it has long been associated to be a cell type of origin for LUAD. By linking dynamic allelic effect pattern with TF-centered regulon activity pattern, we could find an example of LUAD susceptibility gene potentially regulated under cell-state-associated gene regulatory network (GRN). Namely, *SFTPA2*, int-eQTL gene and TWAS gene in AT2 cells, can be potentially regulated through *CEBPD* and *FOXP2*, where *FOXP2* itself is a AT2-specific eQTL. *CEPBD* is part of the CCAAT/enhancer-binding family of transcription factors, and it was reported to play a role in response to acute lung injury and lung development^72,73^. Whereas *FOXP2*, which is a member of the forkhead box family of transcription factors, has been shown to play a role in lung development^74^. *SFTPA2* is a TWAS gene from a locus not reaching genome-wide significance in the GWAS, and our TWAS analysis indicated that higher levels of *SFTPA2* are associated with LUAD risk. Notably, pathogenic germline variants in surfactant genes such as *SFTPA2* and *SFTPA1* have been previously associated with lung cancer and interstitial lung diseases^75,76^. Our results suggested that in response to stress, *CEBPD* can upregulate *SFTPA2* expression in AT2 cells mediating LUAD risk, although the exact mechanism of how higher expression can increase risk still needs to be explored in future studies. Overall, we were able to identify gene programs that can dynamically contribute to differentiation and potentially tumorigenesis.

We recognize that our study has several limitations. First, while our strategy enabled preservation and enrichment of epithelial cells, we did not enrich endothelial and stromal cells, which could reduce biological insights, as these cell types can also play a role in lung cancer etiology, although their contributions may be smaller than epithelial and immune cells. Second, our trajectory analyses focused on gene programs that may be involved in the transition between AT2 to AT1. Although historically the onus has been on AT2, recent studies have implicated that in certain contexts (i.e., in the presence of *KRAS/EGFR/TP53* driver mutation(s)), AT1 may revert to a stem-like state^62^, which warrants future studies focusing on these particular contexts.

Overall, we provided an outline on a disease customizable sc-eQTL dataset. Our dataset increases the diversity in current genomic datasets enabling us and others to identify context-specific susceptibility genes to further our understanding of lung cancer etiology.

## Methods

### Human subjects and tissue collection

Patients’ tissues were obtained under a protocol approved by Yonsei University Health System, Severance Hospital, Institutional Review Board (IRB 4-2019-0447, 4-2022-0706), and informed consent was obtained from each patient prior to surgery. Donors’ consent included secondary use of their samples for research purposes, alongside providing data, which contained personally identifiable information such as sex, age and medical center. All samples were collected at a single hospital (Yonsei University in Seoul, Korea) to primarily study risk factors and prognoses of respiratory diseases; all participants self-reported as ethnically Korean, female, and a never-smoker (smoked < 100 cigarettes in their lifetime). All experiments were performed following protocols compliant with the applicable regulatory guidelines. Normal lung parenchyma tissue samples were collected from 129 patients who underwent curative-intent surgery for pathologically confirmed or radiologically suspected primary lung cancer. Samples were obtained from tumor-distant regions, defined as areas located more than 2 cm away from the tumor edge. The non-malignant status of tumor-distant lung parenchyma was confirmed based on histological evaluation of separately collected tumor-distant lung tissue from the same patients by pathologists. All patients with pathologically confirmed primary lung cancer were diagnosed with lung adenocarcinoma. Most of the patients had early stage of lung adenocarcinoma (79.1% stage IA/IB), while all participating patients were treatment naïve prior to surgery and did not present other active lung diseases, such as interstitial lung disease or infectious lung disease. Four patients underwent surgery for radiologically suspected malignancy but were diagnosed with benign lesions on postoperative pathological examination (nodular dense fibrosis, multifocal interstitial fibrosis, chronic granulomatous inflammation, and lymphoplasmacytic proliferation, respectively). In these patients, tissue resection was performed at sites located more than 2 cm away from the lesion edge. Our sample size was determined based on a power calculation study, which determined > 120 samples provided sufficient power for sc-eQTL mapping^30^. Detailed characteristics of the patients in our study including age, tumor stage, pathology, and smoking status are included in Supplementary Table 1.

### Tissue dissociation and cryopreservation

Cell isolation and cryopreservation from tissues was performed as described in Long et al^29^. In brief, ∼2 cm^3^ size tissues were collected and placed in MACS Tissue Storage Solution (Miltenyi Biotec cat. 130-100-008) within 30 mins of surgical resection and kept at 4°C and processed within 3 hours. Tissues were then manually chopped into approximately 2 mm-diameter pieces and dissociated using Multi Tissue Dissociation Kit 1 (Miltenyi Biotec, cat. 130-110-201) on gentleMACS Octo Dissociator (Miltenyi Biotec, cat. 130-096-427). A reduced (25%) amount of enzyme R was used to create a single-cell suspension. Red blood cells were subsequently removed using Red Blood Cell Lysis Solution (Miltenyi Biotec cat. 130-094-183). Single-cell suspension was frozen using 90% FBS and 10% DMSO and stored in liquid nitrogen until further processing; any transfers were done in the presence of dry ice. Enzymatic digestion and freezing/storage conditions were optimized for epithelial cell retention rate and maximum viability.

### Blood DNA genotyping and imputation

Genotypes of each sample (129 individuals) were assayed from genomic DNA isolated from buffy coat (Qiagen QIAamp DNA blood midi kit #51185) using Illumina Infinium Global Screening Array (v2) at the Cancer Genomics Research Laboratory (CGR) at the National Cancer Institute (NCI). All samples passed sex check and sample contamination verification. These samples were run through a pre-imputation pipeline, which included filtering by sample call rate (<95%), genotype call rate (<95%), HWE p-value < 10^-6^, and MAF < 0.01, and subsequently run through the Will Rayner program (HRC-1000G-check-bim.pl). Namely, A/T & G/C SNPs with MAF > 0.4, or SNPs with more than 20% allele frequency difference compared to 1000 Genome phase 3 or not in 1000 Genome phase 3 reference were further excluded.

Genotypes of the 129 individuals were split into separate chromosomes and subsequently imputed using Michigan Imputation Server and 1000 Genomes (Phase 3 v5, hg19) as a reference panel (EAS). We performed liftover of the imputed genotypes to hg38 using LiftoverVcf. All variant filtering post-imputation was performed using bcftools with criteria set for each analysis.

### Cell type balancing and scRNA-seq library preparation and sequencing

To preserve and enrich for epithelial cells, including cells of lung cancer origin, we utilized FACS sorting and cell type balancing as described in Long et al^29^. In brief, cells were quickly thawed and filtered using 70 μm pre-Separation Filters (Miltenyi Biotec, cat. 130-095-823) to remove debris and subsequently labelled with DAPI (ThermoFisher Scientific, cat. D21490), and antibody markers EPCAM (ThermoFisher Scientific, cat. 25-9326-42), CD31 (BioLegend, cat. 303116), and CD45 (BioLegend, cat. 304006). Live cells (DAPI negative) were then sorted into three groups, “epithelial”: EpCAM+/CD45-, “immune”: EpCAM-/CD45+, and the rest (“endothelial” and “stromal”): EpCAM-/CD45-on BD FACSAria Fusion Flow Cytometer (BD Biosciences). Following a wash (PBS + >= 20% FBS), the cells were mixed at a ratio of 6:3:1 (epithelial:immune:rest) for an individual sample. For each batch, equal numbers of balanced cells from 6 samples (except for 3 batches with 4, 5, or 8 samples) were loaded to 10X Genomics Chromium Controller (100069, 10X Genomics, USA), aiming for ∼20,000 cells or ∼3,333 cells/individual. A total of 23 batches were processed. Library preparation was performed following 10X Chromium Next GEM Single Cell 3’ Reagent Kits v3.1 (CG000315-Rev D, 10X Genomics, USA) protocol. Sequencing was performed by the CGR using Illumina

NovaSeq6000 platform with the following parameters: Read 1, 28 cycles; i7 Index, 10 cycles; i5 Index, 10 cycles; Read 2, 90 cycles.

### Gene expression matrices generation

FASTQ files were generated from raw sequencing data using Cell Ranger (10x Genomics, v7.1.0) ‘mkfastq’ function. FastQC (v0.11.9) was used to check the quality of the sequencing runs. Afterwards, counts matrix were generated by aligning scRNA-seq reads to GRCh38 (hg38) reference genome and quantified using Cell Ranger ‘count’ function.

### Demultiplexing of samples based on genotypes

To demultiplex each batch as well as nominate doublets, we separately ran demuxlet^77^ and vireoSNP^78^ (v0.5.8). For the germline genotype input of both programs, we used cutoffs of variant imputation R^2^ > 0.9 and MAF > 0.05. The balance between the total number and quality of the included variants were determined in a pilot analysis comparing R^2^ > 0.8, 0.9, and 0.95 with an alpha = 0.5 on demuxlet, where R^2^ = 0.9 resulted in the highest number of singlets in an 8-plex setting. Singlet recovery was subsequently compared at this cutoff, and 6-plex was selected with an estimation of ∼80% singlet recovery. For demuxlet, expression matrix was the bam file extracted from cellranger, whereas vireoSNP used this bam file to generate a “cell genotype” using cellsnp-lite (default conditions). vireoSNP was then run with default conditions using cell genotype alongside sample genotype. Demuxlet was run with default conditions with an addition of “—alpha 0 –alpha 0.5” option for better estimations of doublets. Demultiplexing results from both programs were highly concordant, with 99.8% agreement in our final dataset; for the 0.2% that disagreed, we utilized demuxlet assignment.

### Quality control, normalization, and integration

DropletQC^79^ (v0.9) was used to identify “empty” droplets containing ambient RNA. DropletQC utilized ‘possorted_genome_bam.bam’ files generated from Cell Ranger as an input and subsequently calculated nuclear fraction (intronic reads/ (intronic + exonic reads). “Empty” droplets were defined by low nuclear fraction and UMI count. Concurrently, Scrublet^80^ (v0.2.3) was applied to identify doublets based on their expression signatures, where we adjusted the expected doublet rate based on the number of detected cells in each batch. Alongside this, we utilized demuxlet and vireoSNP to identify doublets based on genotypic differences. The results from DropletQC, Scrublet, demuxlet, and vireoSNP were incorporated as metadata in Seurat (v5) object. All identified “empty” droplets and Scrublet doublets were removed, and doublets that were marked by both demuxlet and vireoSNP were also removed. All subsequent analysis was performed in Seurat unless stated otherwise. To identify any remaining low-quality cells and/or outliers (which could be indicative of remaining doublets), we calculated the median and median absolute deviation (MAD) number of features expressed in each sample. Cells that had more or less than 2*MAD boundary from the median number of features were removed. Lastly, high mitochondrial DNA (mtDNA) transcripts are associated with unhealthy or dying cell states. Cells that had more than 20% mtDNA were excluded; this threshold was determined based on the distribution profiles of our data and previous studies of lung tissue^26,29^. In total we removed >40% of our initial cells; this stringent QC is to ensure that we retained only high-quality single cells for our subsequent analyses. A breakdown of number of cells removed in each step is summarized in Supplementary Table 3. Following QC, each batch was normalized with ‘SCTransform’, where the top 2,000 variable genes were identified. Following normalization, we used principal component analysis (PCA) to calculate the top 50 PCs. The batches were then integrated using Harmony to correct for batch effect; we observed that batch effect was largely controlled as most of our individuals were represented in most of our cell types (Supplementary Table 5).

### Clustering and cell type annotation

Initial cell-level automatic annotation was carried out with Azimuth (v0.5.0), using HLCA annotation as a reference to guide the clustering and annotation process. Clustering was conducted using the ‘FindClusters’ function of the Leiden algorithm at a resolution of r = 0.1, and ‘RunUMAP’ for dimensionality reduction and visualization. The resulting clusters were grouped into four cell type categories based on Azimuth annotations and canonical marker gene expression (Epithelial: *EPCAM*, Endothelial: *CLDN5*, Immune: *PTPRC*, Stromal: *COL1A2*). Re-clustering was then performed within each category using resolutions from r = 0.45 to r = 3.60, aiming to produce a similar number of clusters as in the HLCA cell types from the Azimuth analysis. Manual annotation was then performed by comparing differentially expressed genes from each cluster to our curated list of marker genes (Supplementary Table 4). NSForest^31^ (v3.9) was run with Python (v3.8), which identifies the necessary and sufficient genes to distinguish clusters. This analysis was performed within categories and then compared to our gene list to assist annotations. Neighboring clusters with similar expression profiles and annotations were merged, while those showing traits of doublets (e.g., high UMI counts and/or multiple marker gene expressions) were excluded. Manual annotations were cross-checked with Azimuth annotations to identify any cell types obscured within larger clusters. When increasing the resolution and re-clustering alone did not distinguish these obscured cell types, the clusters were separated out, and these smaller groups were re-clustered repeatedly before being reintegrated until clear patterns emerged, allowing for their identification and annotation. The resulting cell annotations from each cell category were mapped onto the initial Seurat object to produce the final annotated dataset and remove unmatched cells. NSForest was run on this final dataset, which includes 41 cell types, to identify distinctive markers for each class. A subset of these NSForest markers matched canonically known markers from HLCA (Results), which was used by Azimuth in initial cell type annotation, showing concordance between the approaches.

### Immunofluorescence Staining of Paraffin-Embedded Human Lung Tissue Sections

Paraffin-embedded human lung tissue sections were deparaffinized and rehydrated through a series of xylene and ethanol washes. To reduce red blood cell-associated autofluorescence, sections were incubated with 3% hydrogen peroxide (H_2_O_2_) for 10 minutes at room temperature. Tissue sections were incubated with primary antibodies against surfactant protein C (SFTPC, Invitrogen, PA5-17680, 1:200 dilution), Uteroglobin/SCGB1A1 (R&D Systems MAB4218, 1:100 dilution), and UGRP1/SCGB3A2 (R&D Systems AF3545, 1:00 dilution) overnight at 4°C. Following primary antibody incubation, the sections were washed and subsequently incubated with appropriate secondary antibodies: FITC-conjugated secondary antibody (Santa Cruz, SC-2359, 1:200), Alexa Fluor 647 (Invitrogen, A48272, 1:200), and Alexa Fluor 594 (Invitrogen, A11058, 1:200) for 1 hour at room temperature. After secondary antibody incubation, tissue sections were washed and mounted with DAPI-containing mounting medium to counterstain nuclei. Immunofluorescence images were acquired using the Axio Scan.Z1 slide scanner (Carl Zeiss). Images were visually assessed and interpreted for our protein of interest using ZEN software (Zeiss).

### Pseudo-bulking and eQTL mapping

For eQTL mapping, we fit a linear model using TensorQTL to estimate the effect of variants within +/-1MB of TSS (where start == end-1) of a gene. For gene expression files, we performed pseudo-bulking for 33 out of 41 cell types, where these 33 cell types had at least 40 individuals with at least 5 cells in that given cell type, a threshold chosen based on previous studies^26,34^. We selected genes that were expressed in at least 10% of all cells in our dataset (8,175 genes, median zeros per gene in each tested cell type, Supplementary Table 13). We then extracted the raw counts for these genes for each of the 33 cell types (gene x cell matrix) and performed sum aggregation to generate gene x samples matrices. We performed normalization using Seurat NormalizeData function, followed by quantile normalization. These gene expression matrices were also used to generate 20 top PEER factors^81^ (PFs) for each cell type to account for latent factors to be used as a covariate to increase sensitivity in regression analysis. For germline genotypes, we used imputation R^2^ > 0.3 and MAF > 0.05 for inclusion in eQTL analysis. To account for population structure, we generated 10 genotype PCs using plink2^79^ for the 129 individuals after pruning highly correlated SNPs (--indep-pairwise 250 50 0.9). Covariates optimization was performed in each cell type to maximize eGene discovery, where the number of PFs used was determined based on the detected eGenes (i.e., minimum PF numbers up to 20 where eGene discovery is plateauing or near-plateauing). eGene discovery was not affected by adding one or more genotype PCs in most cell types. The final covariates used in *cis*-eQTL mapping were age, 3 genotype PCs, and a customized number of PFs for each cell type (median = 14, range = 2-20, Supplementary Table 6). Permutation scheme (beta approximation) was used to adjust p-value for multiple-testing correction, and Storey’s method was used for FDR correction for eGene declaration in each cell type (q-value < 0.05).

Alongside TensorQTL, we also performed eQTL mapping using negative binomial regression (jaxQTL)^82^, to potentially boost power and reduce false positive rates as it may better model the sparse count data in smaller cell types. The input was the same as TensorQTL except that jaxQTL uses raw pseudo-bulked expression counts. Overall, jaxQTL did not meaningfully boost power in our dataset, as the numbers of detected eGenes were similar to those with TensorQTL in 27 of 33 cell types, including smaller cell types (Supplementary Fig. 15). Thus, we used the results from TensorQTL for our eQTL findings and the subsequent analyses.

### Functional annotation of eQTL SNPs

For functional annotation of the detected eQTLs, we first performed LD pruning (--indep-pairwise 250 50 0.9) of 6,309,736 SNPs that were used for eQTL mapping down to 99,784 SNPs. Among them, we collapsed eQTL SNPs (genome-wide significant eQTLs) from each cell type to make a “pseudo-lung” eQTL SNPs. We used the LD-pruned non-eQTL SNPs as the background. For functional annotation of these SNPs, we incorporated ENCODE ChromHMM 15 state annotation (E96, Lung) and used annotate_regions function (annotatr (v3.22)) to get the amount of overlap between the SNPs and these annotated regions. We then performed Fisher’s exact test to gauge depletion or enrichment (odds ratio) of the pseudo-lung eQTL SNPs versus the background SNPs. FDR correction (Benjamini-Hochberg) was performed on the extracted p-values, and significance was defined as FDR < 0.05.

### Harmonization of eQTL effect for cell type sharing

Mashr^38^ was utilized to correct effect size estimates between cell types to gauge cell-type-specificity and sharing. Top eQTLs for all cell types (4,372) were used as the “strong subset” to generate data-driven covariance matrices, and a subset of 10,000 random eQTLs were used to learn the correlation structure. Using these, we fit the full model to compute posterior summaries (betas, standard errors, and lfsr). We subset the results in each cell type to only consider the 4,372 top eQTLs, and significance was defined as lfsr < 0.05. Patterns of cell-type-specificity or cell type/category sharing was evaluated by taking significant eQTLs (lfsr < 0.05) in each cell type and iteratively comparing it to those from the other cell types to obtain the shared proportion. Lastly, to evaluate effect size sharing of significant eQTLs between cell types, we considered an eQTL effect as shared with another cell type when it is in the same allelic direction and between 0.5 and 2 folds.

### Cell-type-specific eQTL SNP enrichment and integration of snATACseq data

To gauge functional enrichment of eQTL SNPs in the context of gene promoters and distal enhancers, we took the cell-type-specific and shared eQTL SNPs based on mashr lfsr < 0.05 and mapped their distances to the TSS of their linked genes. These TSS distances and the linked genes were extracted from TensorQTL outputs and graphed for cell-type-specific eQTL SNPs and those shared across two or more cell categories. For cell type aware functional annotation of eQTL SNPs, we incorporated snATACseq data using similarly processed lung tissues from our single-nucleus multiome dataset^29^. Cell-type-specific and shared ATAC peaks were extracted from the multiome dataset for 22 matching cell types. In the scenario where there is not an exact match, the closest cell type was chosen; for example, “Lymphatic EC mature” in our dataset was matched to “Lymphatic” in the multiome dataset. Note that our cell type marker genes largely overlap those used in the multiome study with additional distinctions of sub-cell types. We then asked how many of our cell-type-specific or shared eQTL SNPs overlapped respective cell-type-specific ATAC peaks, shared peaks across two or more categories, or any other peaks.

### Colocalization and fine-mapping

eQTL colocalization analysis was performed using coloc^40^ (v.5.2.3) to nominate susceptibility genes from lung cancer GWAS loci. We used the default settings of coloc which applies the approximate Bayes factor (ABF) colocalization hypothesis under a single causal variant assumption (default). We used summary statistics from two lung cancer GWAS, Byun et al^13^. (multi-ancestry populations, 42 loci in total combining total lung cancer, LUAD, LUSC, and SCLC subtype analyses) and Shi et al^14^. (EAS population, 30 loci, LUAD-only). For each locus, we searched within the +/-100 KB window of GWAS top SNPs and selected the loci that contained at least one eQTL SNP, which left us with 40 loci from Byun et al and 25 loci from Shi et al study. We then performed colocalization analyses using all overlapped variants in the defined loci between lung cancer GWAS and eQTL summary statistics in each cell type.

Posterior probability of H4 (where two traits are associated with the same causal variant) (PP.H4) > 0.7 was used to demarcate high confident colocalized signals. To help prioritize CCVs and identify functional variants for the 6q22.1 (*ROS1*) and 10q25.2 (*TCF7L2*) locus, fine-mapping was performed using *susie*^59^ within the coloc package. We performed the analyses using both our eQTL and East Asian LUAD GWAS (Shi et al) summary statistics. In-sample LD matrices for fine-mapping were generated using genotypes of 4,544 control individuals from the Female Lung Cancer Consortium in Asia (FLCCA) from the Shi et al GWAS dataset as well as 129 individuals from the eQTL dataset. Genotypes (hg19) were filtered following pre-imputation steps similar to eQTL genotype QC and subsequently imputed using TOPMed Imputation Server and TOPMed (R3 hg38) as the reference, which concurrently performed liftover to hg38. The genotypes were then filtered to only include common and high confidence variants (MAF > 0.05 and R^2^ > 0.5), which were used to generate an LD matrix using plink (v1.9). *susie*-nominated credible set(s) (CS) information from GWAS and eQTL datasets were used for variant prioritization for functional validation.

### TWAS

TWAS analysis was conducted using FUSION suite of tools^42^. Expression weights for each cell type were calculated using the pseudo-bulk expression matrices and genotypes that were used for eQTL mapping. This is done by first estimating heritability for each gene with each cell type using GCTA-GREML. We then generated prediction models of each gene based on the *cis-*SNPs (±1MB from TSS) using four models (best linear unbiased prediction, least absolute shrinkage and selection operator or LASSO, elastic-net, and top SNPs) with cross-validation performed for each model. Genes with heritability (p-value < 0.05) and cross-validation (p-value < 0.05) were retained for TWAS analyses. Using these gene weights, expression was imputed to Shi et al LUAD GWAS SNPs, and the in-sample LD matrix from the 4,544 FLCCA genotypes (described above) was used to account for relationships between SNPs. FDR correction was performed on TWAS p-values to account for multiple testing, FDR < 0.05 was used to nominate LUAD susceptibility genes within each cell type.

### Cell-state interacting eQTL and GRN analyses

To investigate whether eQTLs exhibit dynamic behavior across alveolar cell states, we applied a single-cell negative binomial mixed-effects (NBME) model. For each gene, UMI counts were modeled as a function of genotype and its interaction with pseudotime, while adjusting for sample-level covariates (age), cell-level fixed effects (library size [nUMI] as offset, percentage of mitochondrial UMIs), the top six expression principal components (expPCs), and the top three genotype PCs (gPCs). expPCs were generated from normalized (Seurat NormalizeData) single-cell expressions data for each tested cell type using the PCAforQTL^83^ package with default conditions. A random effect was included to account for donor-specific intercept.

Pseudo-time values were inferred using Monocle3^48,49^, a widely used and effective tool for trajectory analysis. The trajectory plot from Monocle3 indicated that cells in partition 1 (the largest partition) bifurcated into two branches. To focus on a biologically relevant subset, we selected the branch containing cell types of AT2, Alveolar Transitional Cells, and AT1, and applied NBME only in these cells. To capture dynamic effects, pseudo-time was divided into six quantiles (cells with infinite pseudo-time were removed), with the ordinal values (1–6) treated as a continuous variable in the NBME model. Dynamic eQTLs were identified with an FDR threshold of <0.05. To further localize effects along the trajectory, we examined each pseudotime quantile separately by modeling the relationship between UMI counts and genotype, adjusting for the same covariates as above. These effect estimates were used for fitting a linear and/or quadratic model and plotting allelic effects. To identify potential transcription factors that may be mediating allelic effects, we performed motifbreakR^56^ analysis on the int-eQTL SNP alongside correlated SNPs with threshold of p < 1e-04 to denote SNPs that may alter transcription factor binding sites.

Gene regulatory network [GRN] analysis was carried out using *SCENIC* (pyScenic)^55^. First, we extracted expression and meta data for the three cell types of interest (AT2, Alveolar Transitional, and AT1) and generated a loom file. Correlation structures between gene expression across cell types were used to nominate transcription factors (TFs) and their target genes to generate a “regulon”. These regulons were then refined by pruning indirect targets (i.e., genes that do not have an enrichment for corresponding motifs of the TF in <10kb from the TSS). Regulons (TFs and their target genes) activity scores are available on a cell-by-cell basis.

To incorporate trajectory, we extracted pseudo-time values from our trajectory analysis and matched them to their corresponding cell, separating them into the same 6 partitions of our NBME analysis. To gauge dynamic regulon activity, we fitted regulon activity as a function of quantile pseudo-time for each regulon, using a linear and/or quadratic model, and significance was determined at p-value < 0.05. To gauge which pathways these dynamic regulons were functioning in, we performed GO enrichment (clusterProfiler) with dynamic TFs and their corresponding dynamic eQTL genes.

### CRISPRi

Monoclonal A549-dCas9-ZIM3 cell line was established in our previous study^58^ We established H1975-dCas9-ZIM3 polyclonal line in this study. First, plasmid pRC0528_Lenti-dCas9-ZIM3-Blast (encoding dCas9-ZIM3, from Dr. Raj Chari) was co-transfected into HEK293T cells with psPAX2, pMD2-G, and pCAG4-RTR2 packaging vectors to generate lentivirus particles.

Lentivirus particles were collected at 72 h after transfection and filtered (0.45 micron) to remove dead cells. The virus titer was measured by using RETRO-TEK HIV-1 p24 Antigen ELISA Kit (ZeptoMetrix). H1975 cells were infected with dCas9-ZIM3 lentivirus by adding polybrene at concentration of 8ug/mL, then selected for 10 days with 8 mg/mL blasticidin for the generation of an H1975-dCas9-ZIM3 polyclonal stable cell line.

CCVs prioritization criteria were detailed in the Results. In short, we incorporated our previously published MPRA dataset alongside ENCODE functional annotation from multiple lung related datasets to nominate likely functional variants. For CRISPRi, non-targeting and adeno-associated virus site 1 (AAVS1) sgRNAs were used as negative controls. The controls alongside sgRNAs targeting the selected CCVs were designed by Raj Chari, Ph.D. from Frederick National Laboratory for Cancer Research, who also provided us the plasmids (Backbone, gRNA sequence are in Supplementary Table 14). Plasmid purification was performed using QIAGEN Midi kit (#12145) from an overnight culture (100 mL of LB with 100 ug/mL of Carbenicillin), and the quality and quantity of plasmids were verified via gel electrophoresis (0.7% agarose) and Nano-drop. Lentivirus for gRNAs were generated using HEK293T cells using the same procedure described above and collected after 72 hours. Viral titer was measured using Takara’s Lenti-X GoStix Plus (#631281). H1975-dCas9-ZIM3 cells for *ROS1* and A549-dCas9-ZIM3 cells for *TCF7L2* were cultured in RPMI media with 10% FBS, 1% antibiotic-antimycotic) (Corning, #30-0004-CI), and 2.5 and 4 ug/mL of blasticidin for H1975 and A549, respectively. Cells were infected with an equal titer of virus (gRNAs) across conditions at 60% confluency. Puromycin was added to their normal media for selection, 1.6 ug/mL and 2.5 ug/mL for H1975-dCas9-ZIM3 and A549-dCas9-ZIM3, respectively. Six replicates were used for each experiment and each virus; three total independent experiments were performed. Cells were collected 72 hours post infection and resuspended in RLT buffer with 1% beta-mercaptoethanol (BME) for RNA isolation using Qiagen RNeasy kit (#74104). Subsequently, cDNA was generated using SuperScriptTM III Reverse Transcriptase kit (#18080051, H1975/*ROS1*) or ThermoFisher High-Capacity cDNA Reverse Transcription kit (#4368813, A549/*TCF7L2*). TaqMan probes (ThermoFisher Scientific) for *ROS1* (Hs00177228_m1), *TCF7L2* (Hs01009044_m1), and *GAPDH* (Hs02786624_g1) were used for qPCR, where three technical replicates were performed and averaged for each of the six replicates from three independent experiments. Cell lines authentication was done using STR testing and are confirmed to be mycoplasma free via Lonza MycoAlert kit.

### In-silico knockdown and pathway analysis

For *in-silico* knockdown of *ROS1* and *TCF7L*2 in single-cell expression data, we extracted raw counts from AT2 cells and filtered out lowly expressed genes (expressed < 10% of all cells) and performed pseudo-bulking (sum aggregate to individuals) followed by normalization (Seurat NormalizeData) them. We stratified individuals by how lowly (25^th^ percentile) or highly (75^th^ percentile) expressed our gene of interest (*ROS1* or *TCF7L2*). We then performed differential gene expression analysis on raw count data using DESeq2 comparing the low vs high groups (32 vs 32). Gene set enrichment analysis (GSEA) was subsequently performed using clusterProfiler, and the top 5 enriched gene sets were plotted.

### Cell growth assays (xCELLigence)

shRNAs constructs targeting TCF7L2 CDS(s)/UTR and control (Supplementary Table 15) were purchased from Sigma. Plasmid purification and lenti-virus generation followed the same procedure outlined in our CRISPRi experiments. A549 cells were cultured in RPMI media (1% antibiotic-antimycotic, 10% FBS) and were infected with an equal titer of virus (shRNAs) across conditions at 60% confluency. Puromycin (5 µg/mL) was added to the culture medium 24 hours after infection, and survived polyclonal cells after treatment with puromycin for 72 hr were cultured in medium with 2µg/ml of puromycin and used for growth assays (xCELLigence).

TCF7L2 knockdown was confirmed by western blotting for TCF7L2 (rabbit monoclonal, Cell Signaling #2569S, 1:1000) and GAPDH (mouse monoclonal, Santa Cruz Biotechnology #47724, 1:500).

For xCELLigence, cells were seeded into RTCA E-plates (Agilent, 5469830001) at 5000 cells per well with 4 technical replicates per condition. They were grown for approximately 5 days with readings (Cell Index, impendence) taken about every 15 minutes. To measure the effect of TCF7L2 knockdown on cell growth, we took a similar approach to a previous study^84^ and applied linear mixed models on the impendence data. The different condition was considered a fixed effect while the technical replicate was considered a random effect. Effect size (beta coefficients) was estimated using the linear mixed-effects function (maximum-likelihood procedure) with R package nlme (v3.1-168). This was calculated on the growth phase of the assay, which we estimated to be from 25-75 hours.

## Statistical analysis

Besides TensorQTL which was ran using R (v4.2), all other analyses using R were performed with R (v4.4). All analyses using Python was performed using v3.8. GraphPad Prism (v10) was used to analyzed CRISPRi data.

## Data availability

The raw sequencing data alongside the final Seurat object and genotype data will be made available on GEO and dbGaP, respectively. Gene expression alongside eQTL results across lung cell types can be searched on a webtool, ISOLUTION: https://appshare.cancer.gov/ISOLUTION/.

## Code availability

The codes used to analyze our dataset can be found on Github (https://github.com/NCI-ChoiLab/Lung_single_cell_eQTL/).

## Acknowledgement

This research was supported in part by the Intramural Research Program of the National Institutes of Health (NIH). The contributions of the NIH author(s) are considered Works of the United States Government. The findings and conclusions presented in this paper are those of the author(s) and do not necessarily reflect the views of the NIH or the U.S. Department of Health and Human Services. The work that is reported was partially supported by grant U19CA203654 (CIA, YH, JB). We thank the helpful advice and support of Dr. Pradeep Dagur at the NHLBI Flow Cytometry Core, Kristine Jones and colleagues at the NCI Cancer Genomics Research Laboratory, and Dr. Raj Chari at the NCI Genome Modification Core. We also thank Dr. Linh Bui-Raborn for helpful discussions. This work utilized the Biowulf cluster computing system at the NIH.

## Titles of Supplementary Tables

Baseline characteristics of tumor-distant normal lung tissues collected in this study

Overview of scRNA-seq QC data after Cell Ranger analysis

Number of cells after filtering likely droplets, doublets, and low-quality

Cell type annotation based on canonical markers

Overview of number of cell types and cells per individual

Overview of cell types used for eQTL mapping

Summary of eGenes and significant eQTLs

Summary of colocalization results using EAS and multi-ancestry summary statistics

Summary of TWAS results using EAS summary statistics

Summary of lung cancer susceptibility genes identified in this study

Overview of significant dynamic eQTLs results

Beta approximation from xCELLigenece

Number of zero counts after filtering lowly expressed genes

Details of gRNA plasmids used for CRISPRi experiments

Details of shRNA plasmids used for cell growth assays

---

**Supplemental Figure 1.**
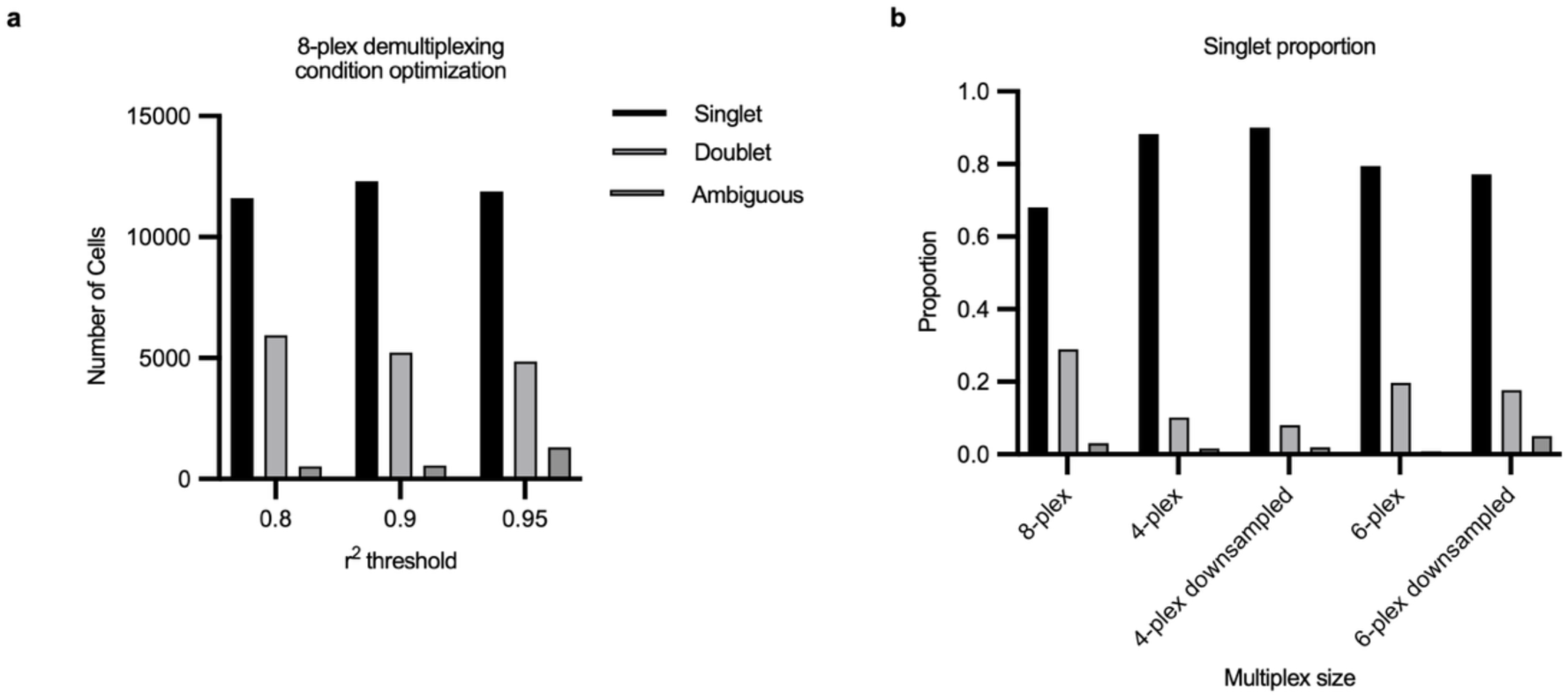
Multiplexing optimization. Sample multiplexing batch size was determined based on the efficiency of the genotype-based demultiplexing to achieve the maximum recovery of cells assigned as singlets and reduce the cost. Germline genotype data from matched blood samples was imputed using Michigan imputation server and 1000 genomes EAS populations as a reference. The imputed genotypes were then filtered by quality (imputation r^2^ > 0.8 to 0.95) and utilized by Demuxlet to match the genotypes from scRNA-sequencing of each cell to assign samples to cells. **a,** Genotype imputation quality cutoffs were compared between r^2^ > 0.8, 0.9, and 0.95 with an alpha = 0.5 in an 8-plex setting. The cutoff with the maximum singlet cells (r^2^ = 0.9) was chosen. Bar ordered follows legend. **b,** 8-plex vs 4-plex were compared for singlet recovery at r^2^ > 0.9 and alpha = 0.5. To account for sequencing depth differences, we also included a down-sampled 4-plex with a matching read-depth to the 8-plex sample. Considering the singlet proportion and sequencing cost, we chose 6-plex, aiming to achieve ∼80% singlet (between 8-plex and 4-plex). A representative 6-plex sample alongside the down-sampled one matching the 8-plex is shown.

**Supplemental Figure 2.**
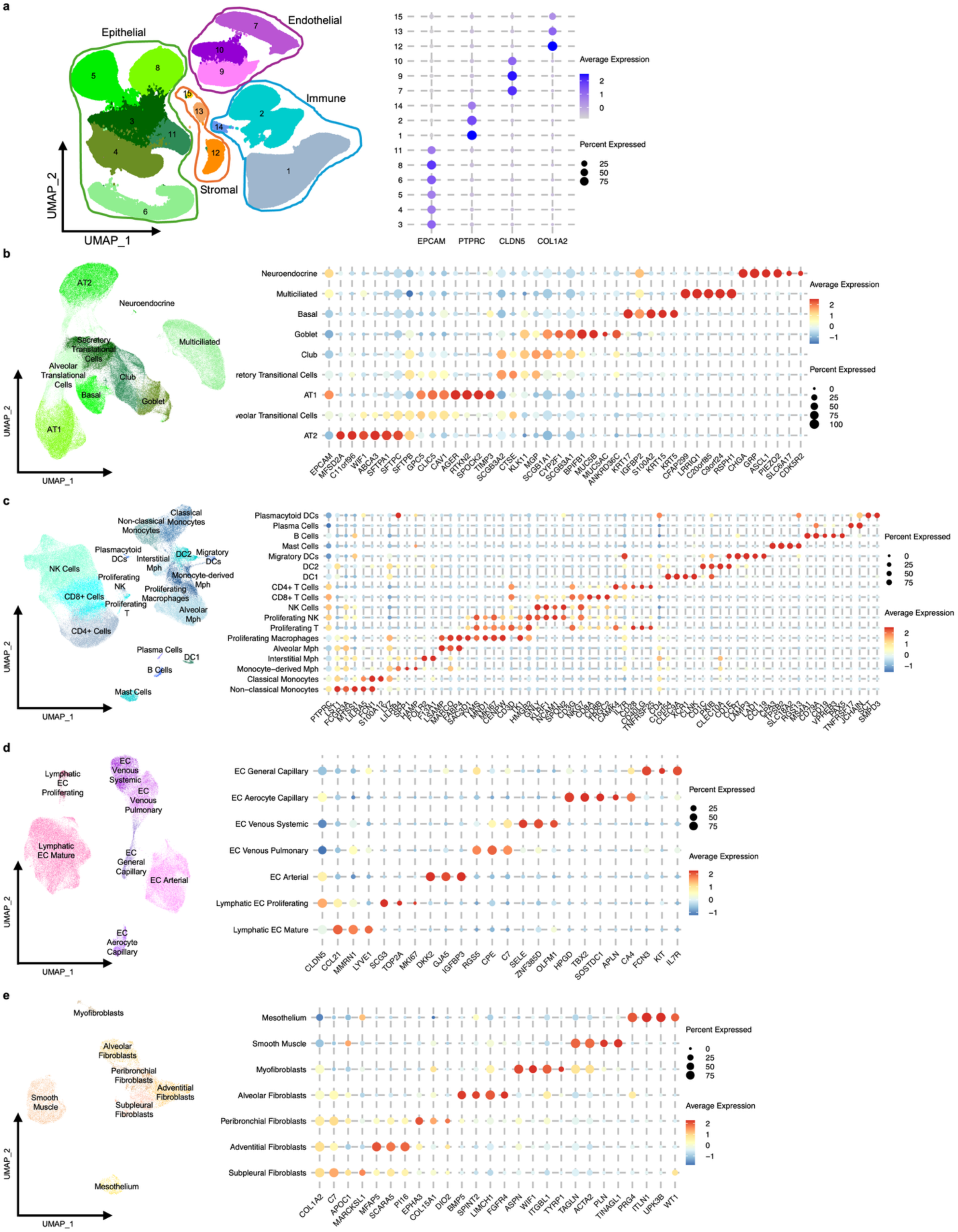
Iterative clustering and cell type annotation in four lung cell categories**. a,** Initial UMAP representation of the whole dataset. Expression of the four canonical markers for epithelial (*EPCAM*), immune (*PTPRC*), endothelial (*CLDN5*), and stromal (*COL1A2*) groups delineate the four cell categories. **b-e**, UMAP plots depicting re-clustering within cell categories: epithelial **b**, immune **c**, endothelial, **d**, stromal. **e,** Expressions of selected marker genes used for cell type annotation are shown on the right panels.

**Supplemental Figure 3.**
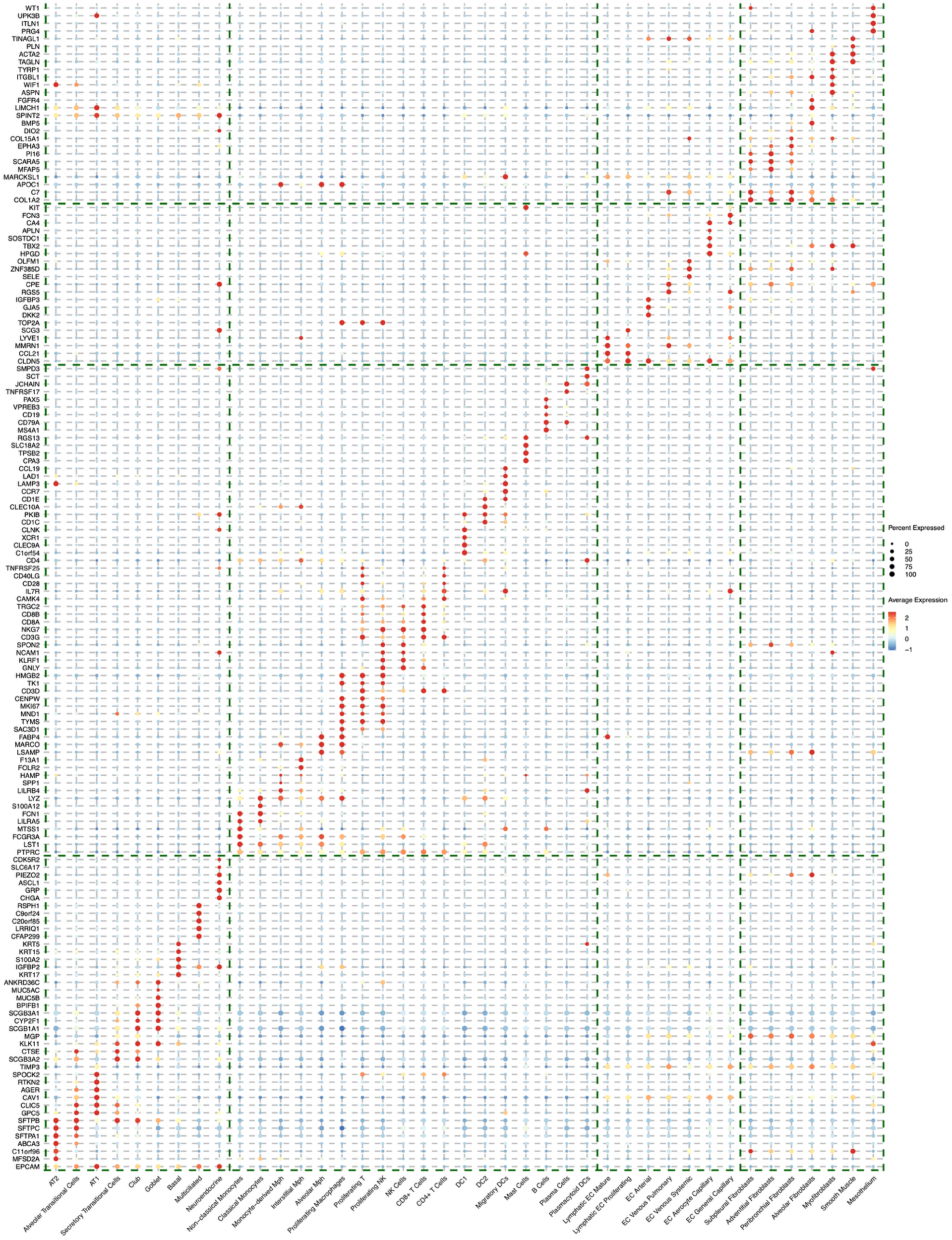
Marker genes for cell type annotation Dot plot showing expression of the selected final makers used to represent the 41 cell types. In this case, markers are on y-axis, while cell types on x-axis. Diagonal expression pattern shows that final set of marker genes are specific to that said cell type.

**Supplemental Figure 4.**
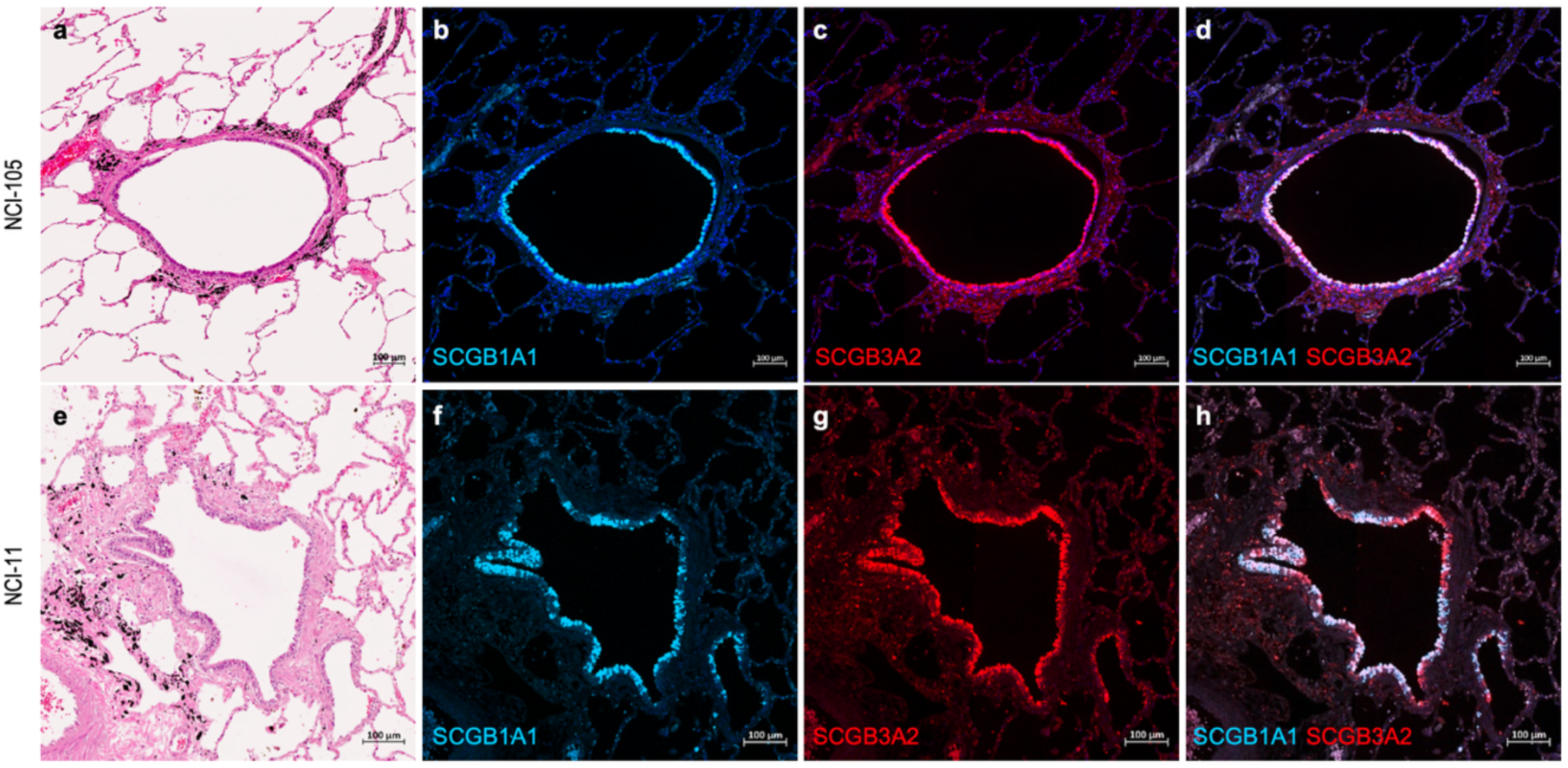
Immunohistochemistry of secretory transitional cells Shown are representative staining results for SCGB3A1 (cyan) and SCGB3A2 (red), with nuclear counterstaining (blue). **a-d** Lung tissue from a patient (NCI-105) exhibiting the lowest abundance of secretory-transitional epithelial cells, as defined by single-cell state signatures. **a,** H&E; **b,** SCGB1A1; **c,** SCGB3A2; **d,** merged. **e-h** Representative staining from a patient (NCI-11) with the highest secretory-transitional cell proportion. **e,** H&E; **f,** SCGB1A1; **g,** SCGB3A2; **h,** merged. In the merged panels, white signals along the bronchial epithelium largely reflect normal club cells. In contrast, regions that appear predominantly red represent segments with diminished SCGB1A1 signal rather than heightened SCGB3A2 levels, consistent with secretory-transitional epithelial populations. The representative patient with high secretory-transitional cell proportions displayed more extensive red-dominant segments compared with the representative low-score individual. Scale bars, 100 µm.

**Supplemental Figure 5.**
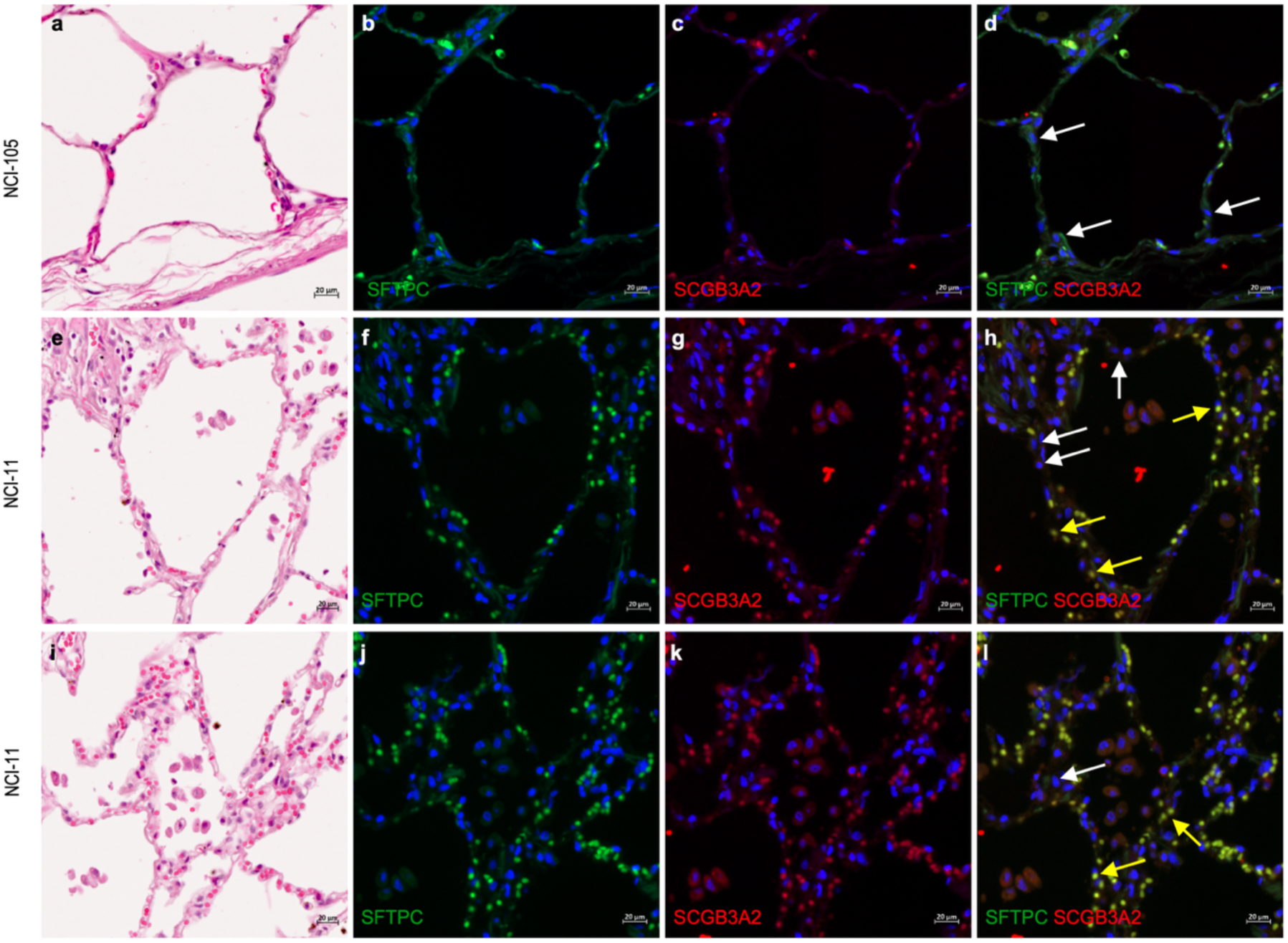
Immunohistochemistry of alveolar transitional cells Shown are representative staining results for SFTPC (green) and SCGB3A2 (red), with nuclear counterstaining (blue). **a-d** Lung tissue from a patient (NCI-105) exhibiting the lowest alveolar transitional cell proportion displays mostly normal alveolar epithelium. **e-h** and **i-l** correspond to two distinct regions sampled from a patient (NCI-11) with the highest alveolar transitional cell proportion, where transitional alveolar epithelial populations are detected. In the merged panels, white arrows indicate representative normal alveolar type II cells characterized by SFTPC signal without detectable SCGB3A2. In contrast, yellow arrows point to representative alveolar type II epithelial cells showing preserved SFTPC with visible SCGB3A2 signal, presumed to represent transitional alveolar epithelial populations. Scale bars, 20 µm.

**Supplemental Figure 6.**
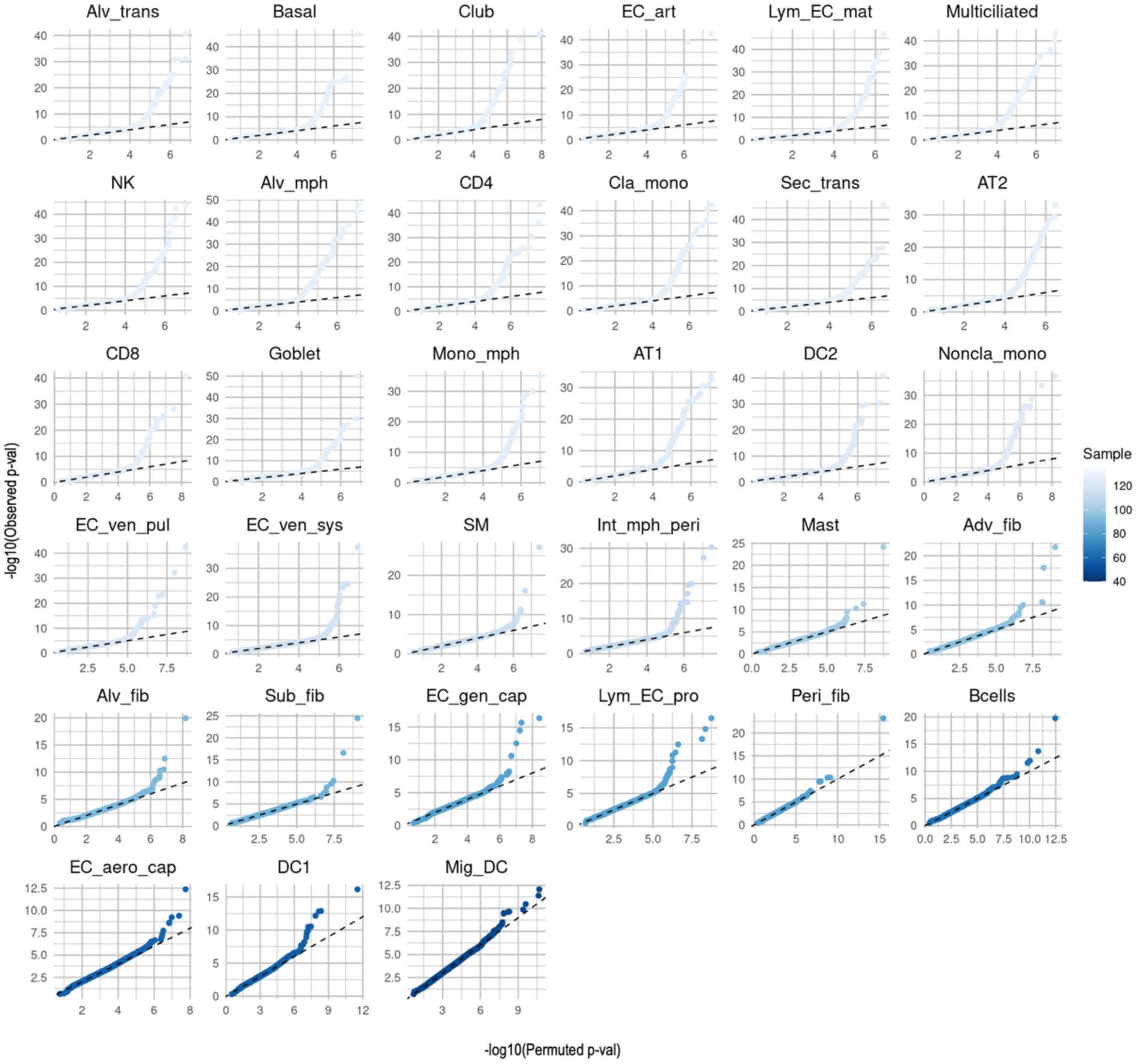
Type I error control of eQTL detection Quantile-quantile plots for each cell-type showing the observed eQTL p-values for top hit per gene (y-axis) against the permutation-based (phenotypes were randomly shuffled for each cell-type) p-values for the top hit per gene (x-axis). Shown on-log_10_ scale. Scale showing the number of individuals with >= 5 cells in each cell type.

**Supplemental Figure 7.**
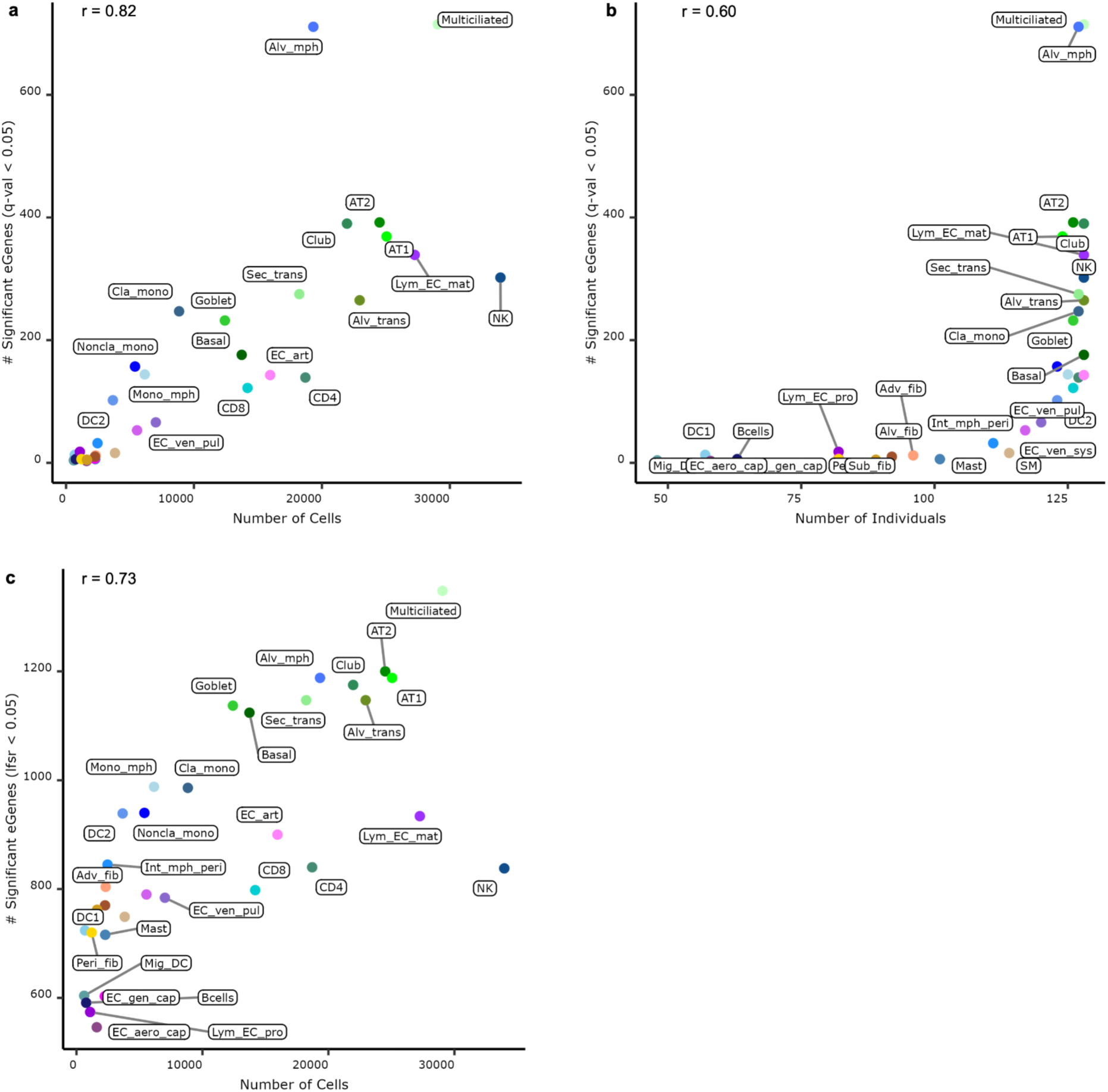
eGene detection power and number of cells or individuals in a cell type Correlation of detected eGenes (q-value < 0.05) with numbers of cells, **a,** and number of individuals, **b,** across the cell types. **c,** Correlation of detected eGenes (q-value < 0.05) with numbers of cells after eQTL effect size harmonization using mashr. Effect size harmonization improved detection power, especially for smaller-size cell types.

**Supplemental Figure 8.**
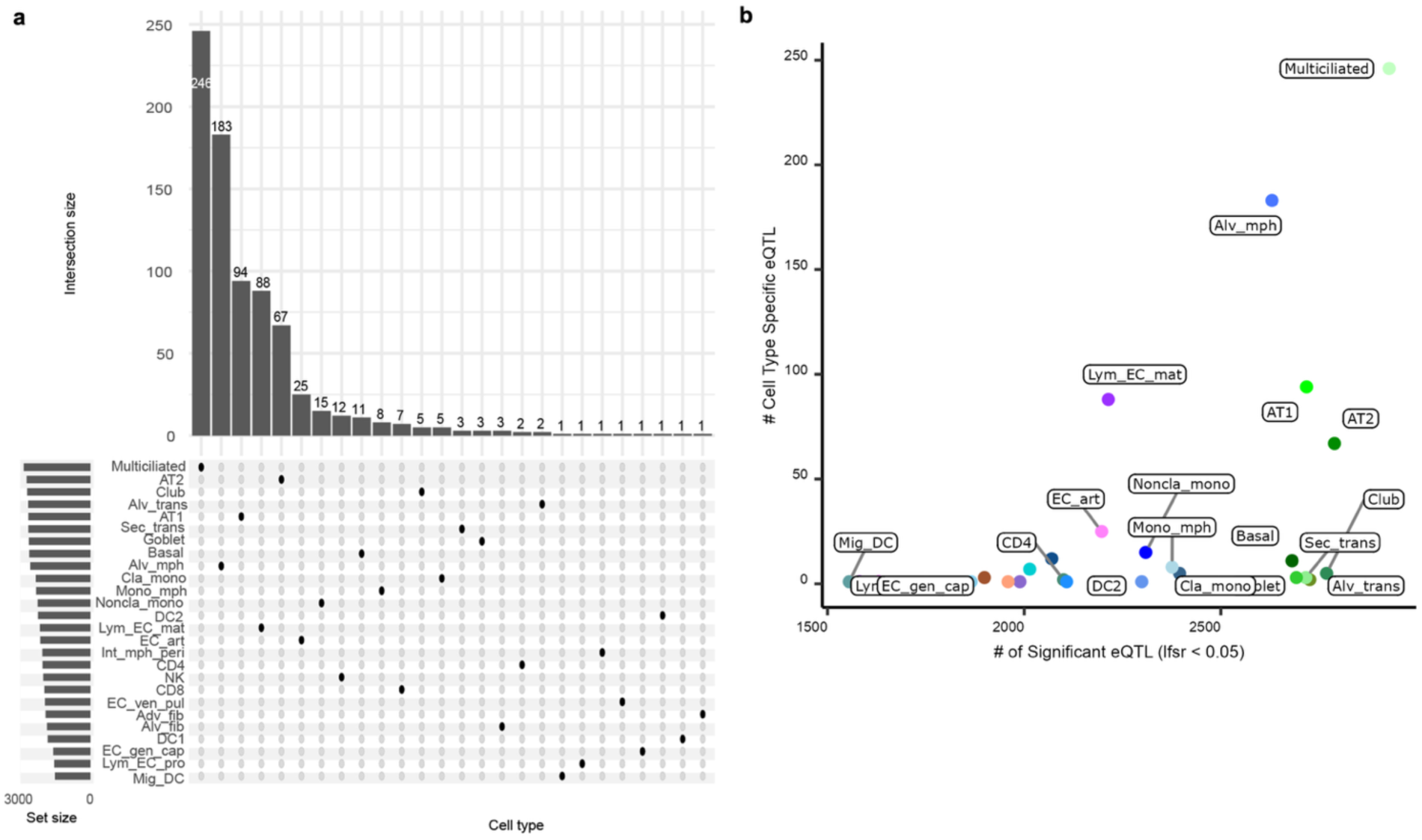
Cell-type-specificity of eQTLs **a,** Upset plot showing the number of cell type specific eQTLs after mashr harmonization. **b,** Scatterplot showing the number of significant eQTLs (lfsr < 0.05) vs number of cell type specific eQTLs on y-axis.

**Supplemental Figure 9.**
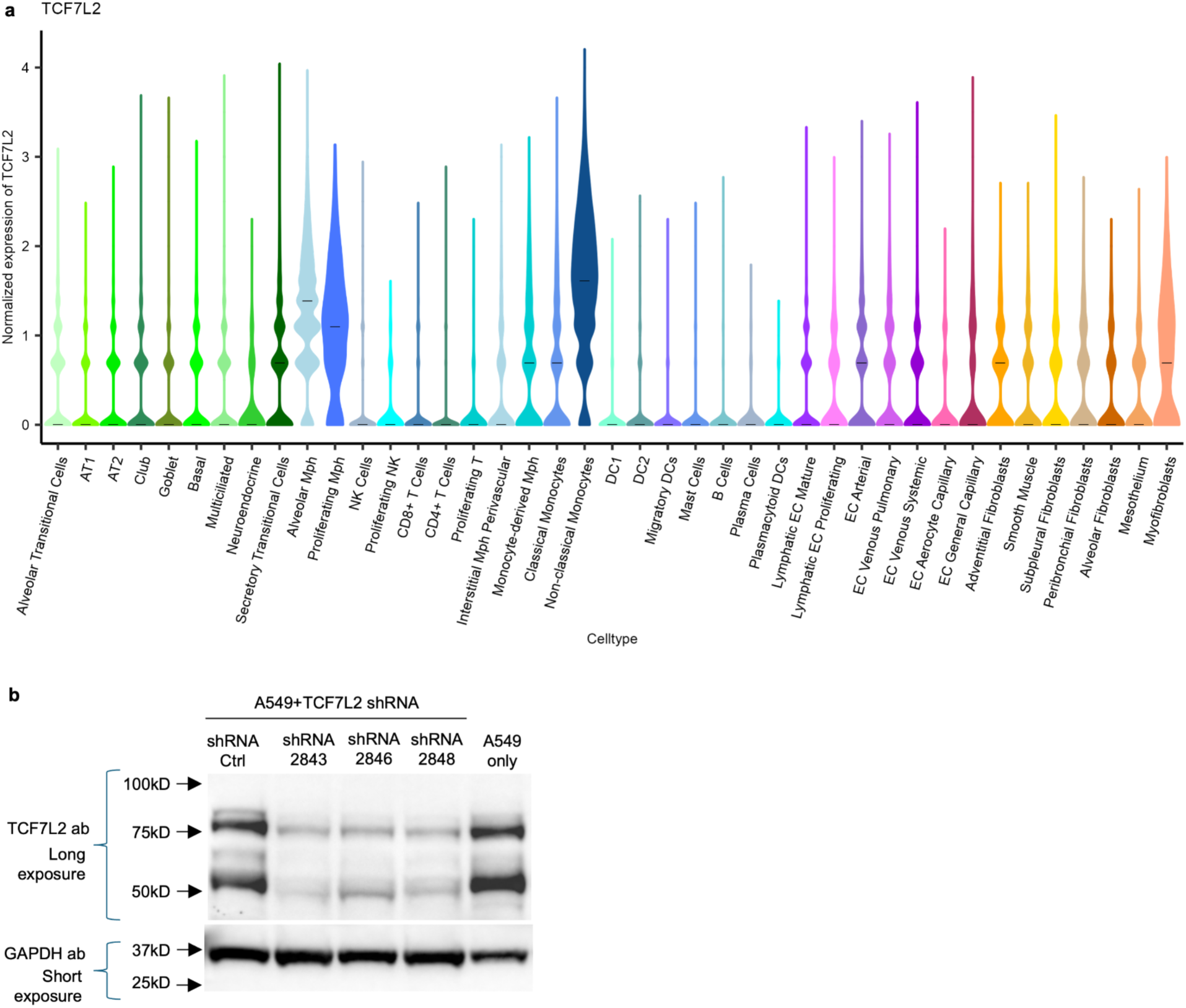
T*C*F7L2 levels in cell types and cell line **a,** Normalized expression of *TCF7L2* across cell types are shown as violin plots. The center line denotes the median, while the violin width reflects the density. **b,** Western blotting showing levels of TCF7L2 in A549 cell lines with control or TCF7L2-targeting shRNA. GAPDH is used as a loading control.

**Supplemental Figure 10.**
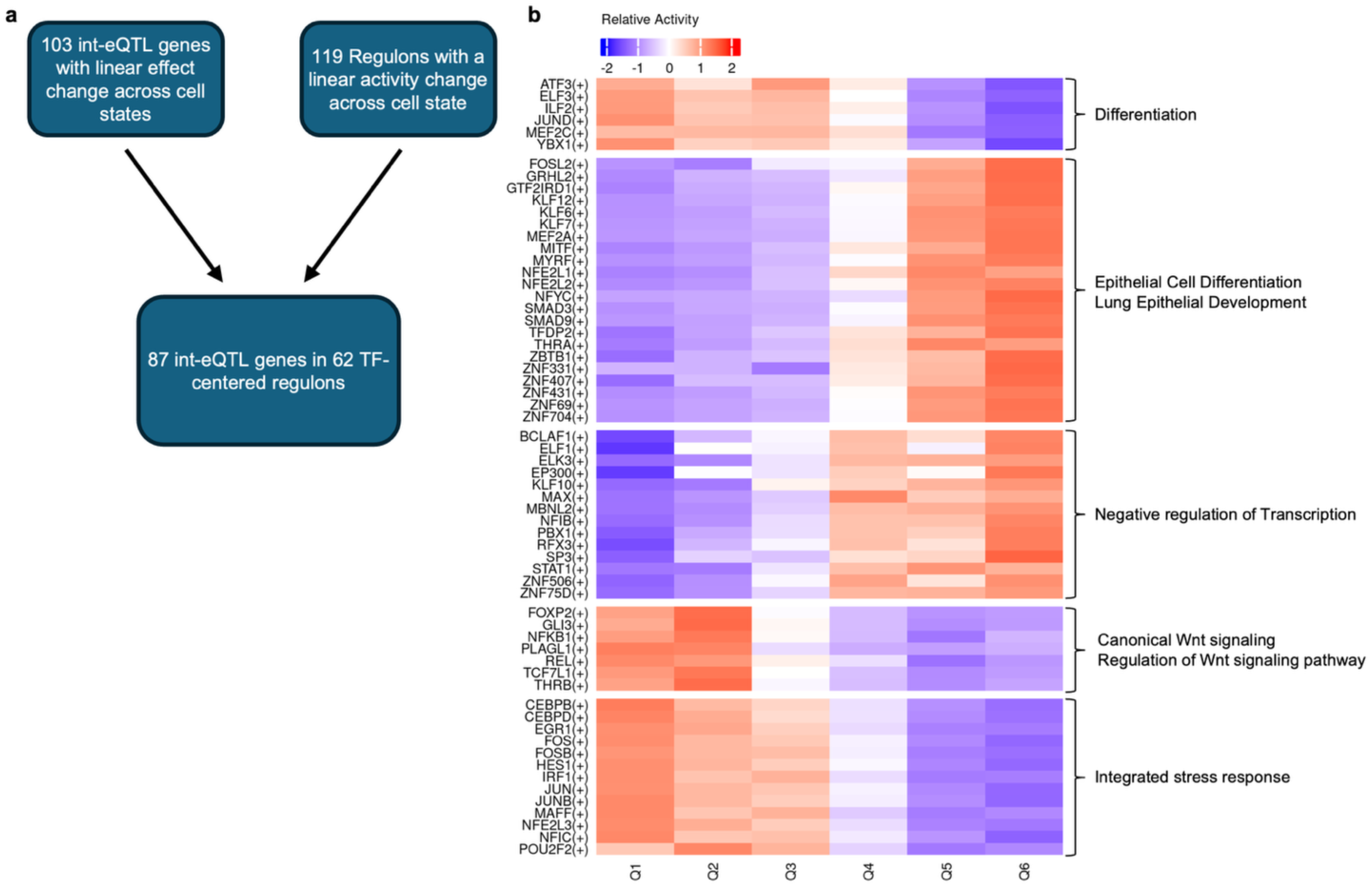
Regulon activity of int-eQTL genes with a linear allelic effect change across cell states **a,** Schematic of overlapping linear int-eQTL genes with linear regulons. **b,** Heatmap of relative activity of the 62 regulons which overlapped the 87 int-eQTL genes, where both regulon activity and int-eQTL effect size fit a linear model across 6 cell states. Genes were clustered into 5 clusters using fuzzy clustering using e1071 package with default settings; TF-centered regulon names shown on the left. Enriched pathways among the TFs and their associated int-eQTLs genes in each cluster are summarized on the right, highlighting overall themes of the top pathways.

**Supplemental Figure 11.**
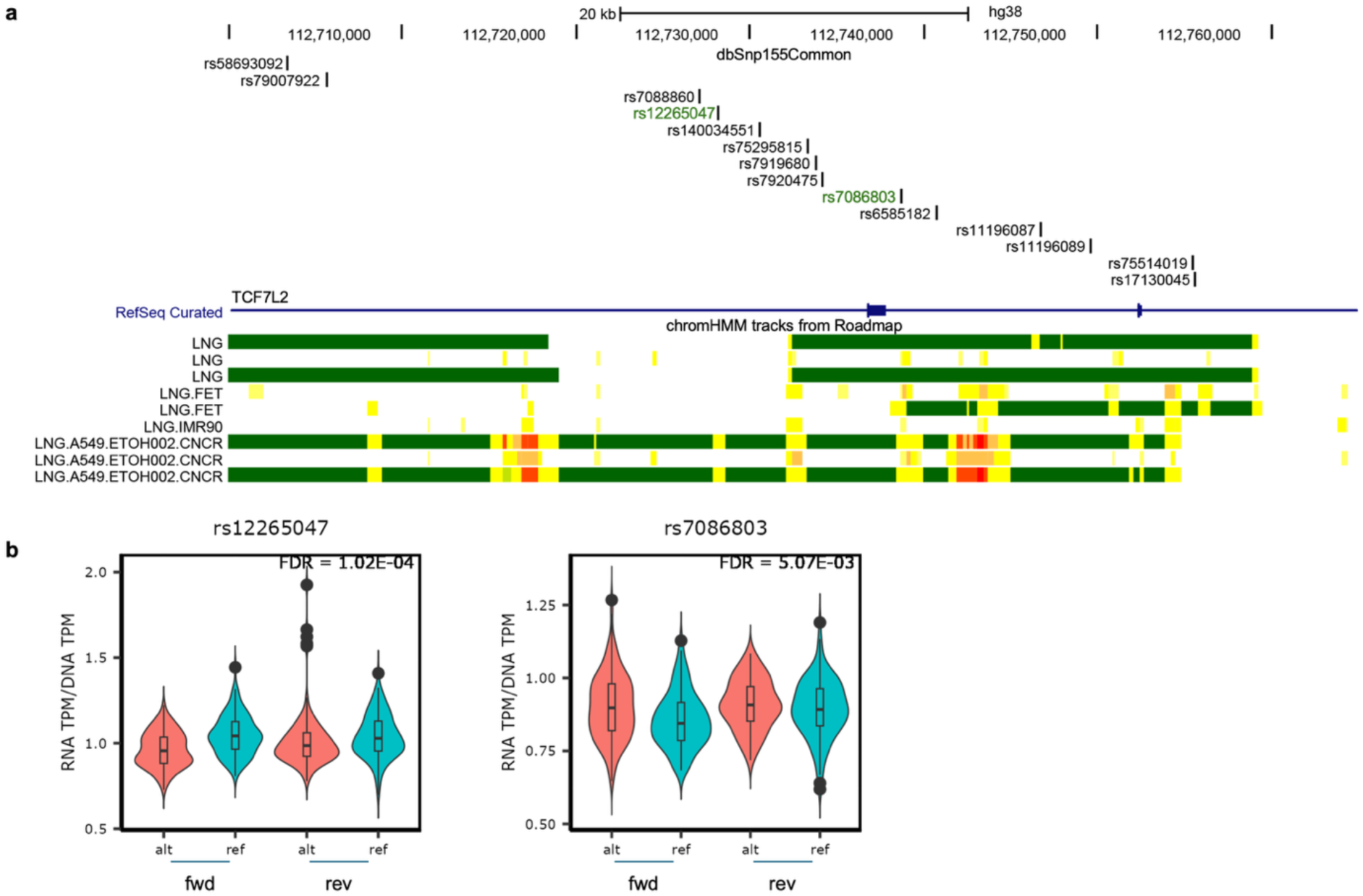
Functional variant(s) nomination for *TCF7L2* **a,** UCSC genome browser image showing 14 candidate causal variants (CCVs) alongside functional annotation from lung-relevant tissues and cell lines. SNPs highlighted in green are the prioritized SNPs based on the overlap with a lung enhancer (yellow shades) or promoter (red shades) and significant MPRA allelic effect in lung cancer cell line(s) (FDR < 0.01). All 14 CCVs show significant MPRA allelic function, and two highlighted variants among them overlap with functional elements in lung. **b,** Allelic effects of rs12265047 and rs7086803 on transcriptional activity (normalized) in H520, shown as violin plots. Center line denotes the median, while the 25^th^ and 75^th^ percentile is marked as the lower and upper line of the box, respectively. Whiskers extend 1.5 times from the 25^th^ and 75^th^ percentiles; outliers are represented as dots. The violin width reflects the density. TPM: Tag-per-million.

**Supplemental Figure 12.**
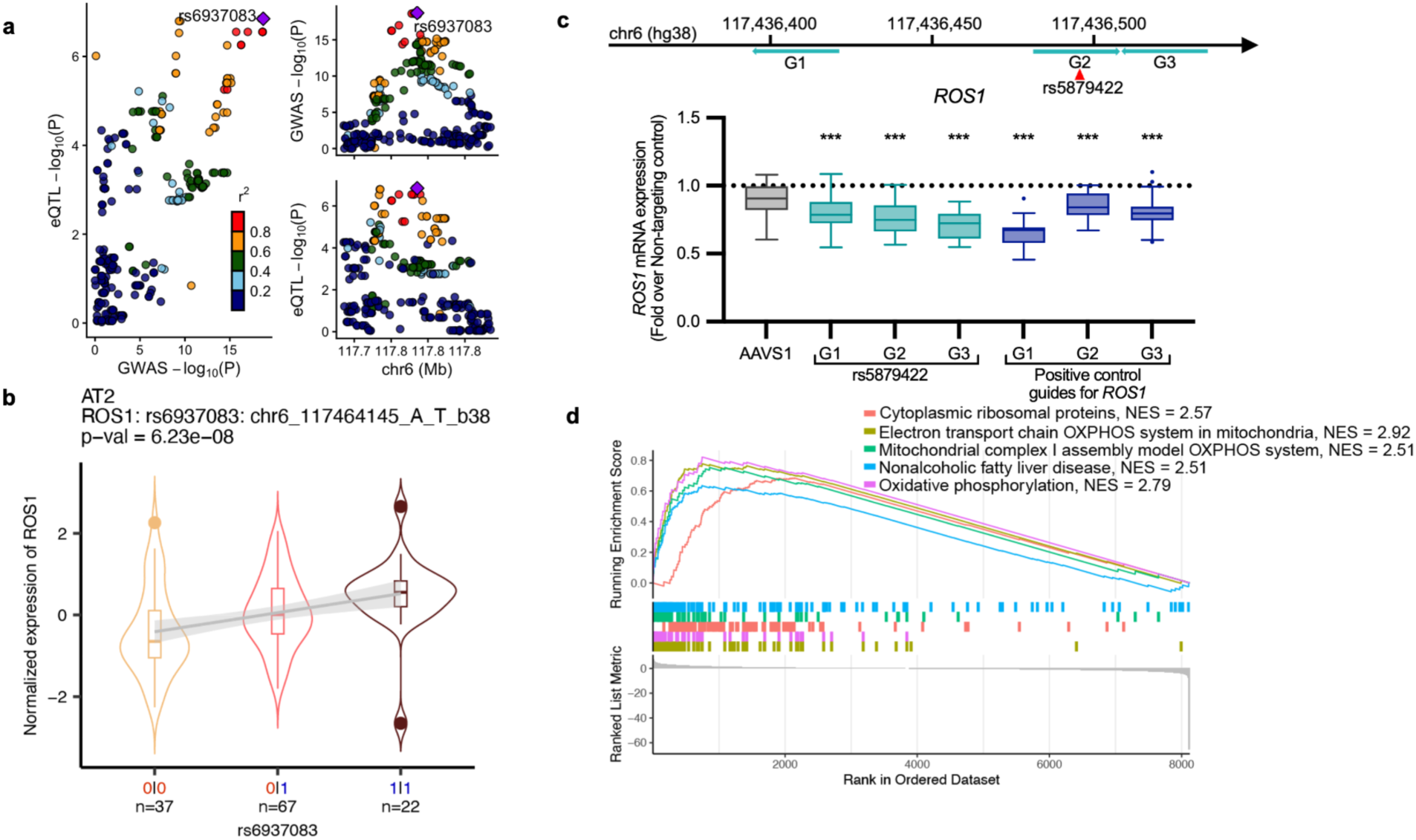
Variant-to-gene validation of *ROS1* **a,** Locus zoom plots of eQTL in AT2 cells for *ROS1* and GWAS (Shi et al EAS, LUAD) p-values of the region (+/-100 kb) around the GWAS top SNP, rs68937083 in the 6q22.1 locus. LD (r2) relationships between the variants are extracted from 1000G EAS Phase 3 v5. **b,** Association between normalized expression of *ROS1* and rs68937083 is shown as violin plots. 0 = reference allele, while 1 = alternative; red indicates risk allele, while blue indicates protective. Center line denotes the median, while the 25^th^ and 75^th^ percentile is marked as the lower and upper line of the box, respectively. Whiskers extend 1.5 times from the 25^th^ and 75^th^ percentiles; outliers are represented as dots. The violin width reflects the density. **c,** Location of guide RNAs targeting the regions around our SNP of interest. Tukey plot shows *GAPDH*-normalized mRNA levels of *ROS1* in H1975 cell line from 6 replicates from three independent experiments (n =18). Fold change of target gene expression over non-targeting control is shown as median with IQR in a box. Whiskers extend 1.5 times IQR, with outliers shown as dots. AAVS1 represents a safe harbor site-targeting gRNA. P-values were calculated using a two-tailed Mann Whitney U test. **d,** GSEA analysis in AT2 cells comparing individuals that lowly (25^th^ percentile) vs highly (75^th^ percentile) expressed *ROS1*. The top 5 most enriched gene sets by NES are shown. NES is normalized enrichment score, where a positive score indicates enrichment of the gene set compared to the reference. *** denotes p-value < 1e-04 while ** p-value < 1e-03.

**Supplemental Figure 13.**
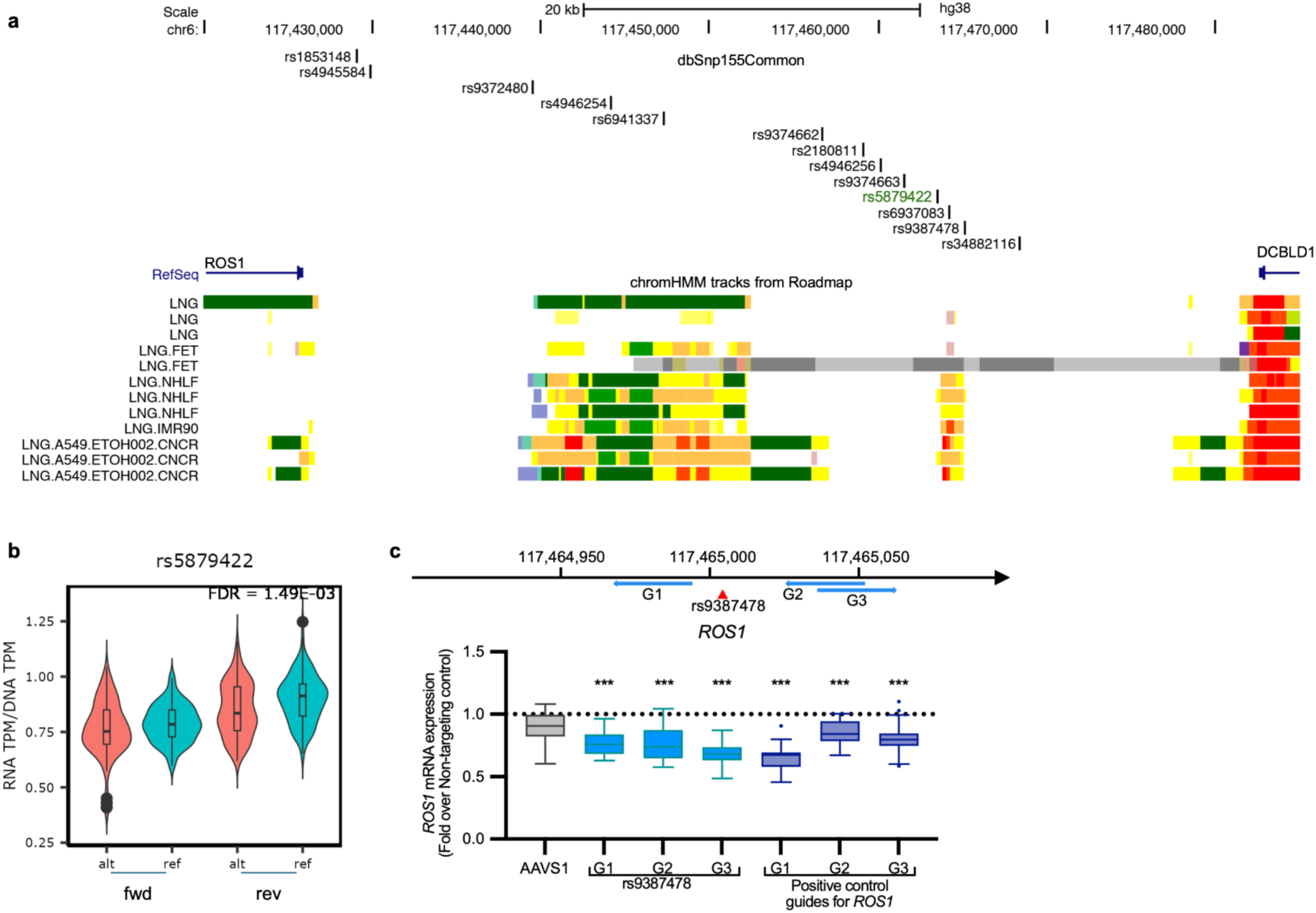
Functional variant(s) nomination for *ROS1* **a,** UCSC genome browser image showing 13 candidate causal variants (CCVs) alongside functional annotation from lung-relevant tissues and cell lines. SNPs highlighted in green are the prioritized SNPs based on the overlap with a lung enhancer (yellow shades) or promoter (red shades) and significant MPRA allelic effect in lung cancer cell line(s) (FDR < 0.01). **b,** Allelic effect of rs5879422 on transcriptional activity (normalized) in A549, shown as violin plots. Center line denotes the median, while the 25^th^ and 75^th^ percentile is marked as the lower and upper line of the box, respectively. Whiskers extend 1.5 times from the 25^th^ and 75^th^ percentiles; outliers are represented as dots. The violin width reflects the density. TPM: Tag-per-million. **c,** Location of guide RNAs targeting the regions around rs9387478, a CCV without MPRA allelic effect but located in the same enhancer element as the prioritized CCV, rs5879422. Tukey plot shows *GAPDH*-normalized mRNA levels of *ROS1* from 6 replicates from three independent experiments (n =18). Median with interquartile range (IQR) is shown; whiskers extend 1.5 times IQR, and outliers are shown as dots. Each dot represents the fold change of *ROS1* over non-targeting control. AAVS1 and positive control guides are replotted from Fig 6c. *** denotes p-vaue < 1e-03.

**Supplemental Figure 14.**
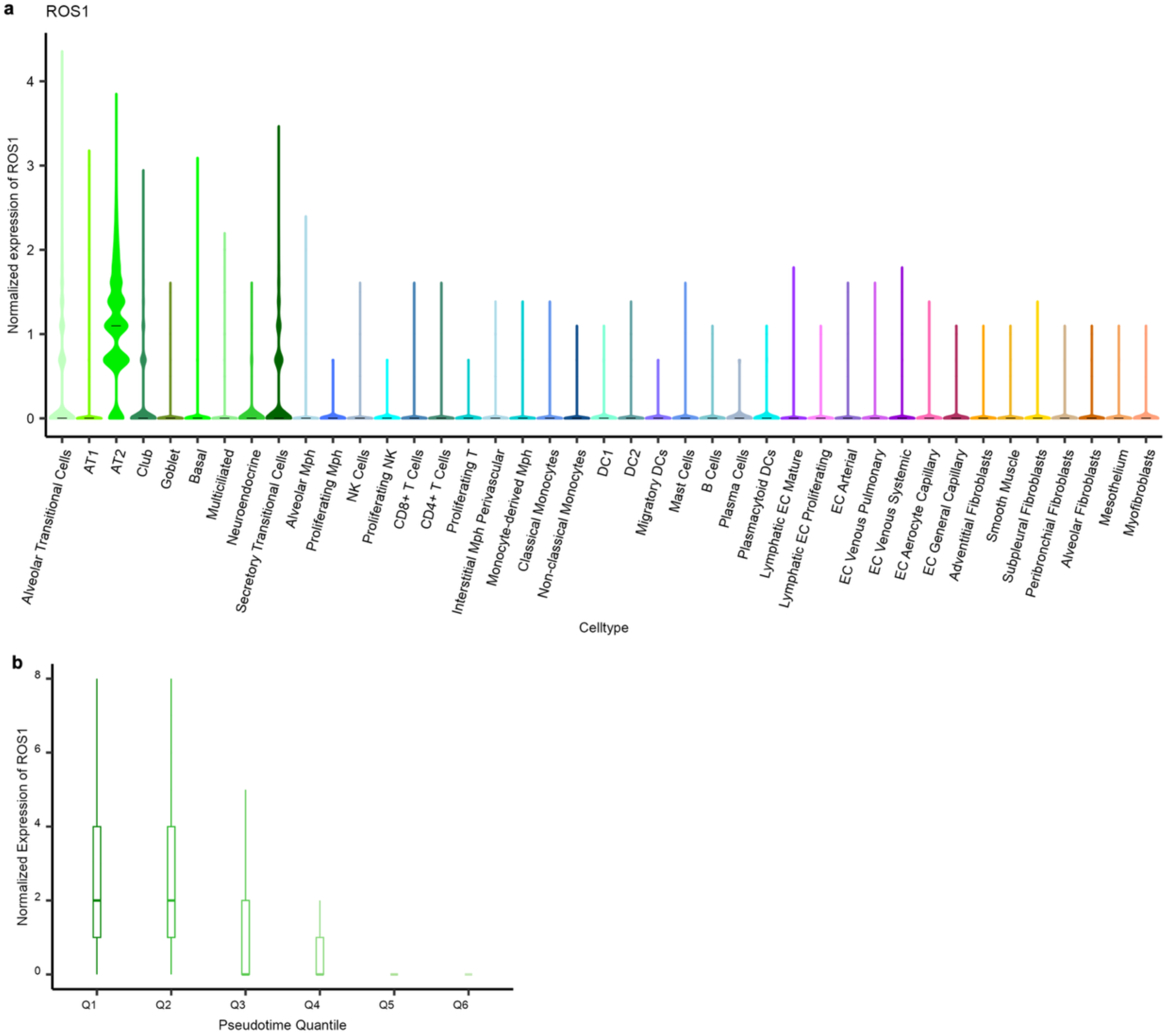
R*O*S1 expression is higher in alveolar progenitor/transitional cell states Normalized expression of *ROS1* across cell types shown as violin plots, **a**, and alveolar epithelial cell states as box plots, **b**, Center line denotes the median, violin width reflects the density. Boxplots show median and the 25^th^ and 75^th^ percentile is marked as the lower and upper line of the box, respectively. Whiskers extend 1.5 times from the 25^th^ and 75^th^ percentiles. Outliers not shown.

**Supplemental Figure 15.**
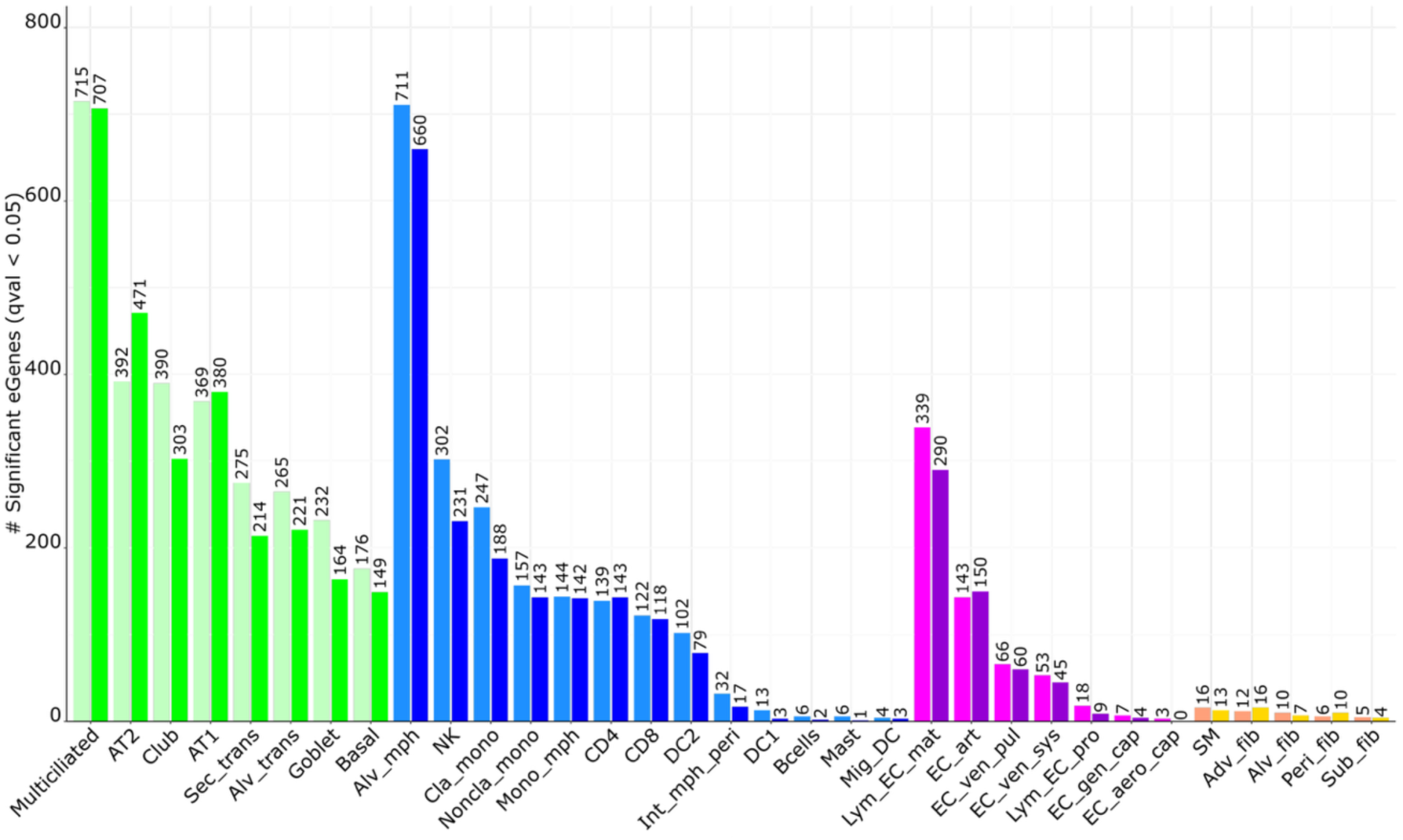
eQTL detection comparisons between linear and negative binomial models Number of declared eGenes (q-value < 0.05) across our 33 tested cell types comparing TensorQTL using linear model (left bars, lighter shades) to jaxQTL using negative binomial model (right bars, darker shades).

## Notes

### Competing Interest Statement

The authors have declared no competing interest.

## References

1. Bray, F. et al. Global cancer statistics 2022: GLOBOCAN estimates of incidence and mortality worldwide for 36 cancers in 185 countries. CA. Cancer J. Clin. 74, 229–263 (2024).

2. Wakelee, H. A. et al. Lung Cancer Incidence in Never Smokers. (2026).

3. LoPiccolo, J., Gusev, A., Christiani, D. C. & Jänne, P. A. Lung cancer in patients who have never smoked — an emerging disease. Nat. Rev. Clin. Oncol. 21, 121–146 (2024).

4. Gomez, S. L. et al. Elevated risk of lung cancer among Asian American women who have never smoked: an emerging cancer disparity. JNCI J. Natl. Cancer Inst. 117, 1104–1109 (2025).

5. Samet, J. M. et al. Lung Cancer in Never Smokers: Clinical Epidemiology and Environmental Risk Factors. Clin. Cancer Res. 15, 5626–5645 (2009).

6. Sampson, J. N. et al. Analysis of Heritability and Shared Heritability Based on Genome-Wide Association Studies for Thirteen Cancer Types. J. Natl. Cancer Inst. 107, djv279 (2015).

7. Mucci, L. A. et al. Familial Risk and Heritability of Cancer Among Twins in Nordic Countries. JAMA 315, 68 (2016).

8. Dai, J. et al. Estimation of heritability for nine common cancers using data from genome-wide association studies in Chinese population. Int. J. Cancer 140, 329–336 (2017).

9. Jiang, X. et al. Shared heritability and functional enrichment across six solid cancers. Nat. Commun. 10, 431 (2019).

10. Byun, J. et al. The Shared Genetic Architectures Between Lung Cancer and Multiple Polygenic Phenotypes in Genome-Wide Association Studies. Cancer Epidemiol. Biomarkers Prev. 30, 1156–1164 (2021).

11. McKay, J. D. Large-scale association analysis identifies new lung cancer susceptibility loci and heterogeneity in genetic susceptibility across histological subtypes. Nat. Genet. (2017).

12. Dai, J. et al. Identification of risk loci and a polygenic risk score for lung cancer: a large-scale prospective cohort study in Chinese populations. Lancet Respir. Med. 7, 881–891 (2019).

13. Byun, J. et al. Cross-ancestry genome-wide meta-analysis of 61,047 cases and 947,237 controls identifies new susceptibility loci contributing to lung cancer. Nat. Genet. 54, 1167–1177 (2022).

14. Shi, J. et al. Genome-wide association study of lung adenocarcinoma in East Asia and comparison with a European population. Nat. Commun. 14, 3043 (2023).

15. Nica, A. C. & Dermitzakis, E. T. Expression quantitative trait loci: present and future. Philos. Trans. R. Soc. B Biol. Sci. 368, 20120362 (2013).

16. Yao, D. W., O’Connor, L. J., Price, A. L. & Gusev, A. Quantifying genetic effects on disease mediated by assayed gene expression levels. Nat. Genet. 52, 626–633 (2020).

17. Bossé, Y. et al. Transcriptome-wide association study reveals candidate causal genes for lung cancer. Int. J. Cancer 146, 1862–1878 (2020).

18. Kim-Hellmuth, S. et al. Cell type–specific genetic regulation of gene expression across human tissues. Science 369, eaaz8528 (2020).

19. Travaglini, K. J. et al. A molecular cell atlas of the human lung from single-cell RNA sequencing. Nature 587, 619–625 (2020).

20. Basil, M. C. et al. Human distal airways contain a multipotent secretory cell that can regenerate alveoli. Nature 604, 120–126 (2022).

21. Kadur Lakshminarasimha Murthy, P., et al. Human distal lung maps and lineage hierarchies reveal a bipotent progenitor. Nature 604, 111–119 (2022).

22. Sikkema, L. et al. An integrated cell atlas of the lung in health and disease. Nat. Med. 29, 1563–1577 (2023).

23. Han, G. et al. An atlas of epithelial cell states and plasticity in lung adenocarcinoma. Nature 627, 656–663 (2024).

24. Yazar, S. et al. Single-cell eQTL mapping identifies cell type–specific genetic control of autoimmune disease. Science 376, (2022).

25. Fujita, M. et al. Cell subtype-specific effects of genetic variation in the Alzheimer’s disease brain. Nat. Genet. 56, 605–614 (2024).

26. Natri, H. M. et al. Cell-type-specific and disease-associated expression quantitative trait loci in the human lung. Nat. Genet. 56, 595–604 (2024).

27. Bian, L. et al. Single-cell eQTL mapping reveals cell-type-specific genes associated with the risk of gastric cancer. Cell Genomics 5, 100812 (2025).

28. Fu, Y. et al. Single-cell eQTL mapping reveals cell-type-specific genetic regulation in lung cancer. Cell Genomics 101100 (2025) doi:10.1016/j.xgen.2025.101100.

29. Long, E. et al. Context-aware single-cell multiomics approach identifies cell-type-specific lung cancer susceptibility genes. Nat. Commun. 15, 7995 (2024).

30. Mandric, I. et al. Optimized design of single-cell RNA sequencing experiments for cell-type-specific eQTL analysis. Nat. Commun. 11, 5504 (2020).

31. Aevermann, B. et al. A machine learning method for the discovery of minimum marker gene combinations for cell type identification from single-cell RNA sequencing. Genome Res. 31, 1767–1780 (2021).

32. Stankovic, B. et al. Immune Cell Composition in Human Non-small Cell Lung Cancer. Front. Immunol. 9, 3101 (2019).

33. Taylor-Weiner, A. et al. Scaling computational genomics to millions of individuals with GPUs. Genome Biol. 20, 228 (2019).

34. Cuomo, A. S. E. et al. Optimizing expression quantitative trait locus mapping workflows for single-cell studies. Genome Biol. 22, (2021).

35. Kim, T.-K. & Shiekhattar, R. Architectural and Functional Commonalities between Enhancers and Promoters. Cell 162, 948–959 (2015).

36. Heinz, S., Romanoski, C. E., Benner, C. & Glass, C. K. The selection and function of cell type-specific enhancers. Nat. Rev. Mol. Cell Biol. 16, 144–154 (2015).

37. Haberle, V. & Stark, A. Eukaryotic core promoters and the functional basis of transcription initiation. Nat. Rev. Mol. Cell Biol. 19, 621–637 (2018).

38. Urbut, S. M., Wang, G., Carbonetto, P. & Stephens, M. Flexible statistical methods for estimating and testing effects in genomic studies with multiple conditions. Nat. Genet. 51, 187–195 (2019).

39. Grandi, F. C., Modi, H., Kampman, L. & Corces, M. R. Chromatin accessibility profiling by ATAC-seq. Nat. Protoc. 17, 1518–1552 (2022).

40. Giambartolomei, C. et al. Bayesian Test for Colocalisation between Pairs of Genetic Association Studies Using Summary Statistics. PLoS Genet. 10, e1004383 (2014).

41. Zhu, M. et al. A cross-tissue transcriptome-wide association study identifies novel susceptibility genes for lung cancer in Chinese populations. Hum. Mol. Genet. 30, 1666–1676 (2021).

42. Gusev, A. et al. Integrative approaches for large-scale transcriptome-wide association studies. Nat. Genet. 48, 245–252 (2016).

43. Bi, H. et al. Clinical characteristics of patients with ROS1 gene rearrangement in non-small cell lung cancer: a meta-analysis. Transl. Cancer Res. 9, 4383–4392 (2020).

44. Drilon, A. et al. ROS1-dependent cancers — biology, diagnostics and therapeutics. Nat. Rev. Clin. Oncol. 18, 35–55 (2021).

45. Wang, Y. et al. SNP rs17079281 decreases lung cancer risk through creating an YY1-binding site to suppress DCBLD1 expression. Oncogene 39, 4092–4102 (2020).

46. Tung, J. J. & Kitajewski, J. Chloride intracellular channel 1 functions in endothelial cell growth and migration. J. Angiogenesis Res. 2, 23 (2010).

47. Liu, K. et al. Tracing the origin of alveolar stem cells in lung repair and regeneration. Cell 187, 2428–2445.e20 (2024).

48. Trapnell, C. et al. The dynamics and regulators of cell fate decisions are revealed by pseudotemporal ordering of single cells. Nat. Biotechnol. 32, 381–386 (2014).

49. Qiu, X. et al. Reversed graph embedding resolves complex single-cell trajectories. Nat. Methods 14, 979–982 (2017).

50. Tian, C. et al. Single-cell RNA sequencing of peripheral blood links cell-type-specific regulation of splicing to autoimmune and inflammatory diseases. Nat. Genet. 56, 2739–2752 (2024).

51. Hancock, D. B. et al. Meta-analyses of genome-wide association studies identify multiple loci associated with pulmonary function. Nat. Genet. 42, 45–52 (2010).

52. Cho, M. H. et al. Variants in FAM13A are associated with chronic obstructive pulmonary disease. Nat. Genet. 42, 200–202 (2010).

53. Jiang, Z. et al. A Chronic Obstructive Pulmonary Disease Susceptibility Gene, *FAM13A*, Regulates Protein Stability of β-Catenin. Am. J. Respir. Crit. Care Med. 194, 185–197 (2016).

54. Chang, L.-L. et al. AKR1C1 promotes non-small cell lung cancer proliferation via crosstalk between HIF-1α and metabolic reprogramming. Transl. Oncol. 20, 101421 (2022).

55. Aibar, S. et al. SCENIC: single-cell regulatory network inference and clustering. Nat. Methods 14, 1083–1086 (2017).

56. Coetzee, S. G., Coetzee, G. A. & Hazelett, D. J. *motifbreakR*: an R/Bioconductor package for predicting variant effects at transcription factor binding sites. Bioinformatics 31, 3847–3849 (2015).

57. Lan, Q. et al. Genome-wide association analysis identifies new lung cancer susceptibility loci in never-smoking women in Asia. Nat. Genet. 44, 1330–1335 (2012).

58. Long, E. et al. High-throughput characterization of functional variants highlights heterogeneity and polygenicity underlying lung cancer susceptibility. Am. J. Hum. Genet. 111, 1405–1419 (2024).

59. Wang, G., Sarkar, A., Carbonetto, P. & Stephens, M. A Simple New Approach to Variable Selection in Regression, with Application to Genetic Fine Mapping. J. R. Stat. Soc. Ser. B Stat. Methodol. 82, 1273–1300 (2020).

60. Wenzel, J. et al. Loss of the nuclear Wnt pathway effector TCF7L2 promotes migration and invasion of human colorectal cancer cells. Oncogene 39, 3893–3909 (2020).

61. Hanahan, D. Hallmarks of cancer—Then and now, and beyond. Cell S0092867425014989 (2026) doi:10.1016/j.cell.2025.12.049.

62. Juul, N. H. et al. KRAS(G12D) drives lepidic adenocarcinoma through stem-cell reprogramming. Nature 619, 860–867 (2023).

63. COPDGene Investigators et al. Genetic loci associated with chronic obstructive pulmonary disease overlap with loci for lung function and pulmonary fibrosis. Nat. Genet. 49, 426–432 (2017).

64. Cho, M. H., Hobbs, B. D. & Silverman, E. K. Genetics of chronic obstructive pulmonary disease: understanding the pathobiology and heterogeneity of a complex disorder. Lancet Respir. Med. 10, 485–496 (2022).

65. Shrine, N. et al. Multi-ancestry genome-wide association analyses improve resolution of genes and pathways influencing lung function and chronic obstructive pulmonary disease risk. Nat. Genet. 55, 410–422 (2023).

66. Liu, Y. et al. High-Penetrance Rare Variants Underlying Familial Lung Cancer Risk: Insights From Genetic Epidemiology of Lung Cancer Consortium. J. Thorac. Oncol. S1556086425029429 (2025) doi:10.1016/j.jtho.2025.12.004.

67. Zhao, M., Jung, Y., Jiang, Z. & Svensson, K. J. Regulation of Energy Metabolism by Receptor Tyrosine Kinase Ligands. Front. Physiol. 11, 354 (2020).

68. Tuupanen, S. et al. The common colorectal cancer predisposition SNP rs6983267 at chromosome 8q24 confers potential to enhanced Wnt signaling. Nat. Genet. 41, 885–890 (2009).

69. Pomerantz, M. M. et al. The 8q24 cancer risk variant rs6983267 shows long-range interaction with MYC in colorectal cancer. Nat. Genet. 41, 882–884 (2009).

70. Sur, I. K. et al. Mice Lacking a *Myc* Enhancer That Includes Human SNP rs6983267 Are Resistant to Intestinal Tumors. Science 338, 1360–1363 (2012).

71. Long, E., Williams, J., Zhang, H. & Choi, J. An evolving understanding of multiple causal variants underlying genetic association signals. Am. J. Hum. Genet. 112, 741–750 (2025).

72. Breed, D. R., Margraf, L. R., Alcorn, J. L. & Mendelson, C. R. Transcription Factor C/EBPd in Fetal Lung: Developmental Regulation and Effects of Cyclic Adenosine 3*,5*-Monophosphate and Glucocorticoids. 138,.

73. Yan, C. et al. CCAAT/Enhancer-Binding Protein δ Is a Critical Mediator of Lipopolysaccharide-Induced Acute Lung Injury. Am. J. Pathol. 182, 420–430 (2013).

74. Shu, W. et al. Foxp2 and Foxp1 cooperatively regulate lung and esophagus development. Development 134, 1991–2000 (2007).

75. Benusiglio, P. R., Fallet, V., Sanchis-Borja, M., Coulet, F. & Cadranel, J. Lung cancer is also a hereditary disease. Eur. Respir. Rev. 30, 210045 (2021).

76. Ducrot, L. et al. Penetrance of interstitial lung disease and lung cancer in carriers of *SFTPA1* or *SFTPA2* pathogenic variants. ERJ Open Res. 11, 01348–02024 (2025).

77. Kang, H. M. et al. Multiplexed droplet single-cell RNA-sequencing using natural genetic variation. Nat. Biotechnol. 36, 89–94 (2018).

78. Huang, Y., McCarthy, D. J. & Stegle, O. Vireo: Bayesian demultiplexing of pooled single-cell RNA-seq data without genotype reference. Genome Biol. 20, 273 (2019).

79. Muskovic, W. & Powell, J. E. DropletQC: improved identification of empty droplets and damaged cells in single-cell RNA-seq data. Genome Biol. 22, 329 (2021).

80. Wolock, S. L., Lopez, R. & Klein, A. M. Scrublet: Computational Identification of Cell Doublets in Single-Cell Transcriptomic Data. Cell Syst. 8, 281–291.e9 (2019).

81. Stegle, O., Parts, L., Piipari, M., Winn, J. & Durbin, R. Using probabilistic estimation of expression residuals (PEER) to obtain increased power and interpretability of gene expression analyses. Nat. Protoc. 7, 500–507 (2012).

82. Zhang, Z. E., Kim, A., Suboc, N., Mancuso, N. & Gazal, S. Efficient count-based models improve power and robustness for large-scale single-cell eQTL mapping. Preprint at 10.1101/2025.01.18.25320755 (2025).

83. Zhou, H. J., Li, L., Li, Y., Li, W. & Li, J. J. PCA outperforms popular hidden variable inference methods for molecular QTL mapping. Genome Biol. 23, 210 (2022).

84. Florez-Vargas, O. et al. Genetic regulation of TERT splicing affects cancer risk by altering cellular longevity and replicative potential. Nat. Commun. 16, 1676 (2025).

